# Behavioral and brain responses to verbal stimuli reveal transient periods of cognitive integration of external world in all sleep stages

**DOI:** 10.1101/2022.05.04.490484

**Authors:** Başak Türker, Esteban Munoz Musat, Emma Chabani, Alexandrine Fonteix-Galet, Jean-Baptiste Maranci, Nicolas Wattiez, Pierre Pouget, Jacobo Sitt, Lionel Naccache, Isabelle Arnulf, Delphine Oudiette

**Affiliations:** Sorbonne Université, Institut du Cerveau - Paris Brain Institute - ICM, Inserm, CNRS, Paris 75013, France; AP-HP, Hôpital Pitié-Salpêtrière, Service des Pathologies du Sommeil, National Reference Centre for Narcolepsy, Paris 75013, France; AP-HP, Hôpital Pitié-Salpêtrière, Service de Neurophysiologie Clinique, Paris 75013, France; Sorbonne Université, INSERM, UMRS1158 Neurophysiologie Respiratoire Expérimentale et Clinique, Paris 75005, France. Paris 75013, France

## Abstract

Sleep has long been considered as a state of disconnection from the environment, with absent reactivity to external stimuli. Here, we questioned this sleep disconnection dogma by directly investigating behavioral responsiveness in 49 napping subjects (27 with narcolepsy and 22 healthy volunteers) engaged in a lexical decision task. Participants were instructed to frown or smile depending on the stimulus type (words vs pseudo-words). We found accurate behavioral responses, visible via contractions of the corrugator or zygomatic muscles, in all sleep stages in both groups (except slow-wave sleep for healthy volunteers). Stimuli presented during states with high (vs. low) values of neural markers indexing rich cognitive states more often yielded responses. Our findings suggest that transient windows of reactivity to external stimuli exist in all sleep stages, even in healthy individuals. Such windows of reactivity could be used to probe sleep-related mental and cognitive processes in real-time across all sleep stages.

---

Sleep has classically been considered as a time when we are dead to the world, with significantly reduced (or absent) reactivity to external stimuli. However, research in recent years has progressively called into question this assumption^1^. First, congruent evidence from event-related potentials (ERP)^2–4^, fMRI^5^ ,or intracranial recordings^6^ have shown that at least low-level sensory processing is preserved across sleep stages. Several studies indicate that sleepers can even process symbolic stimuli at different cognitive levels of representation, including semantic and decisional stages^7^. Furthermore, learning-related sensory cues presented during sleep positively impact subsequent recall of cue-related material upon awakening^8–10^ and can even influence participants’ behavior (e.g. smoking reduction) a week later^11^. While all of these examples of sensory processing during sleep are thought to occur automatically and unconsciously^4,12^, some studies have shown an incorporation of sensory stimuli into reported dream content^13,14^, suggesting that, at least sometimes, external stimuli could be processed up to conscious stages during sleep. However, the lack of single trial evidence of stimulus integration during sleep complicates the exploration of the neurophysiological basis of this complex and variable phenomenon. Obtaining behavioral responses that serve as real-time indicators of subjective reports could allow us to analyze brain dynamics associated with sensory integration in a trial-by-trial manner.

Potentially because behavioral responses have long been assumed to be possible only during wakefulness, they are either rejected from the analysis^12^ or not collected at all in sleep studies. The rare studies which explicitly attempted to measure behavioral responses in sleeping participants discovered manual behavioral responses during N1 sleep (sleep onset)^4,15,16^, but not in deeper sleep stages. However, the loss of limb muscle tone could mask behavioral responses during consolidated sleep. Because facial muscles are less affected by muscle atonia than the limbs are^17^, they could be more suited for assessing behavioral responsiveness. In addition, eye movements persist during REM sleep and can be used to signal lucidity in people who are aware of dreaming while asleep^14,18^ (lucid dreamers). In a collaborative study, we combined eye movements and facial muscle contractions to show that lucid dreamers could respond to queries sent during their dreams in polysomnographically-verified REM sleep^14^.

In the present work, we capitalized on this research strategy to further question the sleep disconnection dogma and to explore stimuli integration at the behavioral and neurophysiological level. We recruited 27 participants with narcolepsy, - who present excessive daytime sleepiness, a short REM sleep latency, and a high frequency of lucid dreams^19^, and 22 healthy participants. They were explicitly instructed to perform an auditory lexical decision task while napping by frowning or smiling three times depending on the stimulus type (word versus pseudo-word). Facial EMG on corrugator and zygomatic muscles was recorded in addition to usual polysomnography signals.

We discovered that accurate behavioral responses were possible across all sleep stages including non-REM N3 sleep (slow-wave sleep) and REM sleep. Furthermore, and regardless of the group or sleep/wake stages, responsiveness was associated with previously validated electrophysiological markers of high-cognitive processing. Finally, we also found electrophysiological and subjective (post-nap reports) evidence for a conscious processing of external stimuli during lucid REM sleep. Our findings demonstrate that sleepers can transiently process external stimuli at a high-cognitive level and respond to them across all sleep stages. These transient communication windows could be used to interact with any sleeper in real time.

## RESULTS

### Participants can behaviorally respond to auditory stimuli across all sleep stages

In this study, we tested participants’ ability to perceive, understand, and behaviorally respond to auditory verbal stimuli across different sleep stages. We included both participants with narcolepsy (NP, n=27) and healthy participants (HP, n=22). Their sleep/wake stage was continuously monitored by polysomnography (EEG, EOG, EMG). Words and pseudo-words were verbally presented in a pseudo-randomized order during daytime naps; 1-min periods of stimulation (ON periods) alternated with 1-min periods during which no stimuli were presented (OFF periods) (Figure 1A). Participants were instructed to perform a lexical decision task by frowning or smiling three times according to the stimulus type (behavior-stimulus matching counterbalanced across participants), every time they heard a stimulus whether they were awake or asleep. As we previously showed^14^, such behavioral responses are visible to the experimenters thanks to surface EMG sensors measuring corrugator (frowning) and zygomatic (smiling) isometric contractions (see example in Figure 1B). At the end of each nap, participants reported: (i) their mental content during the nap, (ii) whether they had a lucid dream, and (iii) whether they recalled having actively performed the lexical task while sleeping. Each nap was labeled as lucid or non-lucid in function of participants’ post-nap subjective report, with REM sleep trials from these naps being labeled as lucid or non-lucid accordingly. Importantly, participants were also instructed to signal their lucidity (if any) with a “mixed-contraction”, by alternating one corrugator and one zygomatic muscle contraction. These objective dream lucidity signals matched participants’ subjective reports upon awakening (for details see Supplementary Results).

**Figure 1.**
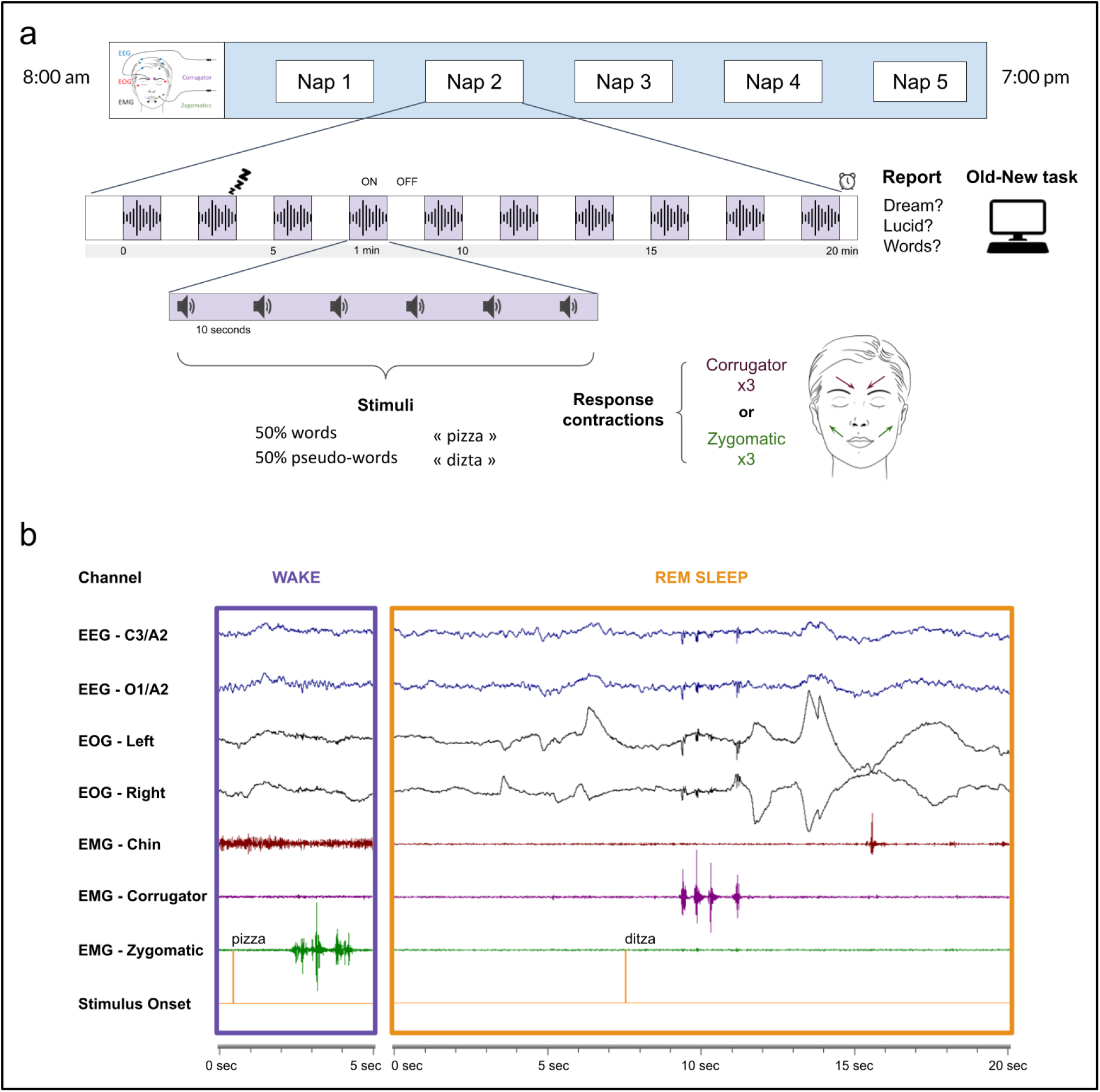
Experimental design for participants with narcolepsy and example of a responsive trial. (a) Participants with narcolepsy went through five 20-minute naps during the same day. In each nap, periods with stimulation (ON) alternated, every minute, with periods when no stimulus was presented (OFF). During the ON periods, participants were presented with words and pseudo-words and asked to either frown (corrugator muscle contractions) or smile three times (zygomatic muscle contractions) in response to the stimuli. Stimuli were presented every 10 seconds (+/- 1 second). Following each nap, participants were asked to report whether: (i) they had any dream; (ii) they were lucid; and (iii) they recalled any words presented during the nap. Immediately after this debriefing, participants performed a forced-choice ‘old/new’ recognition task. Healthy participants went through the exact same procedure except that they had a single 100-min nap. (b) Example of behavioral responses during wake (left, 3 zygomatic muscle contractions in response to a pseudo-word) and REM sleep (right, 4 corrugator muscle contractions in response to a word). The orange vertical line on the last channel indicates the stimulus onset. In this example, we observed the typical markers of REM sleep: low chin tone (EMG), rapid eye movements (EOG), and theta rhythm (EEG).

We first assessed the responsiveness to task-stimuli across sleep stages in the two groups. We compared response rates (corrugator and zygomatic contractions combined) during *ON* and *OFF stimulation periods* (Figure 2A). Importantly, we excluded all responses performed during micro-arousals and only kept periods when participants were asleep according to the sleep scoring rules^20^. As expected, we found significantly higher response rates during ON *vs* OFF periods, both during Wakefulness (HP: 79.8% vs 1.5%, *z* = 31.61, *p* < .0001 after FDR correction; NP: 86.1% vs 2.1%, *z* = 27.02, *p* < .0001) and N1 sleep (HP: 23.2% vs 1.4%, *z* = 11.44, *p* < .0001; NP: 64.2% vs 1.7%, *z* = 18.29, *p* < .0001) in both groups. Crucially, we also found, in both HP and NP, significantly higher response rates in ON *vs* OFF periods during N2 (HP: 4.9% vs 2.1%, *z* = 4.70, *p* < .0001; NP: 20.27% vs 2.2%, *z* = 16.57, *p* < .0001) and (non-lucid) REM sleep (HP: 6.5% vs 2.2%, *z* = 3.59, *p* = .0004; NP: 34.2% vs 1.4%, *z* = 13.93, *p* < .0001). Note that the response rates were higher in NP than in HP. We did not find a significant difference between ON and OFF periods in HP during N3 sleep (0.2% vs 0.1%, *z* = 1.23, *p* = 0.22), but we found significantly more responses during ON than OFF periods in N3 sleep in NP (5.7% vs 2.4%, *z* = 3.31, *p* = .0009). Response rates during ON periods decreased significantly from Wake to N1 sleep, REM sleep, then N2 sleep (in order) in HP. Similarly, they decreased significantly from Wake to N1 sleep, non-lucid REM sleep, N2 sleep, and N3 sleep (in order) in NP (Figure 2A and Table S1). Participants can therefore provide behavioral motor codes with their facial muscles during all sleep stages.

**Figure 2.**
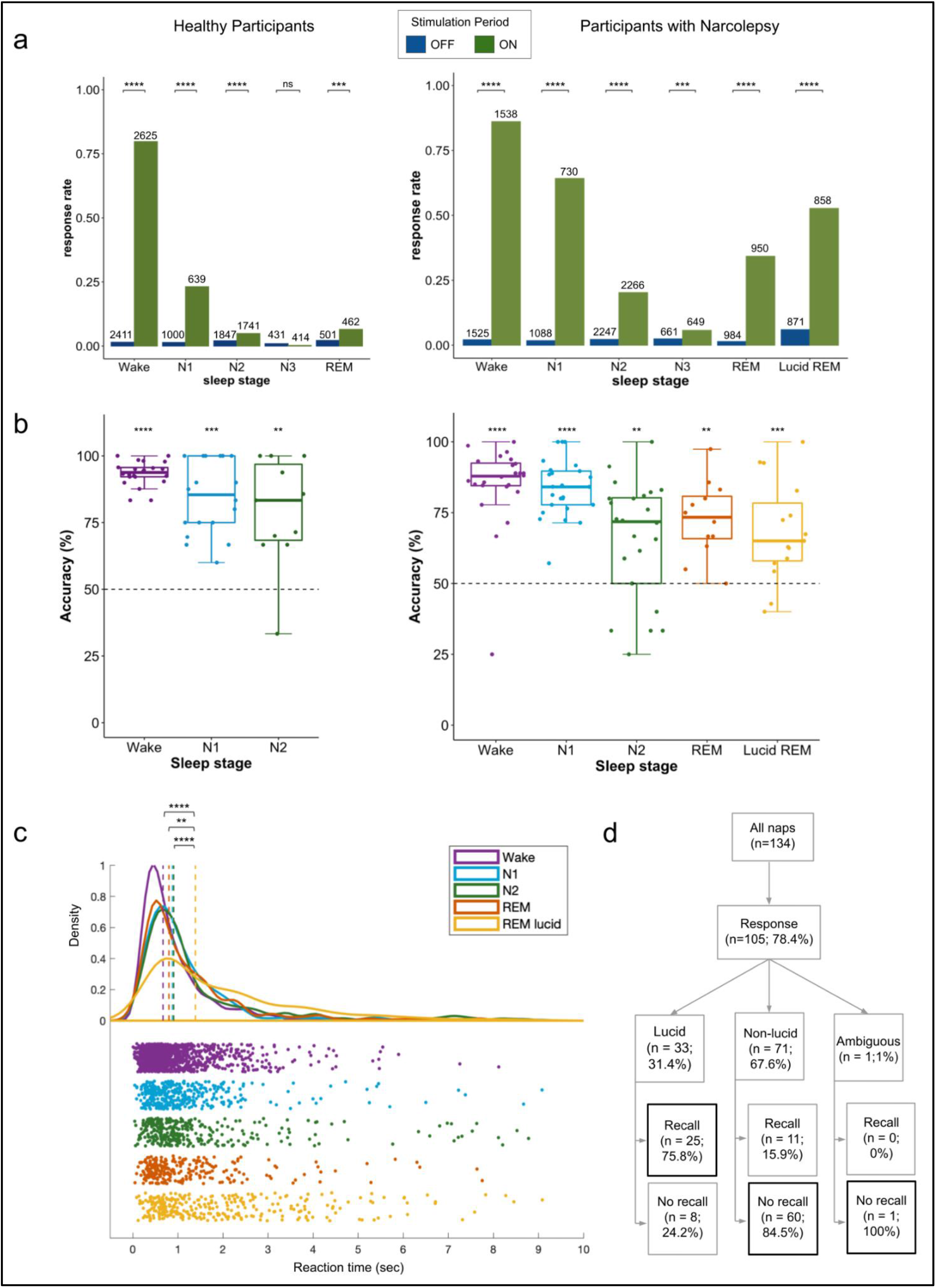
Behavioral results. (a) The overall response rate across different sleep stages during OFF (blue) and ON (green) stimulation periods in participants without (left) and with (right) narcolepsy. The response rate was computed by combining both response types (zygomatic or corrugator muscle contractions), whether the response was correct or not. The total number of trials is indicated on top of the bars. We used binomial generalized mixed-linear models with participants as a random factor for statistical analysis. Significant differences between ON and OFF periods are indicated for each sleep stage. Response rates were significantly larger in ON than in OFF periods in all sleep stages except for N3 sleep in participants without narcolepsy. (b) Accuracy computed over responsive trials in the lexical decision task for participants without narcolepsy -HP (left) and with narcolepsy -NP (right). Only participants with at least 3 responses were included in this analysis. Each dot represents a participant and dashed lines indicate the 50% chance level. Both HP and NP were significantly more accurate than chance in all sleep stages (c) Distribution of reaction times from stimulus onset to response in correct trials (words and pseudo-words) in NP across sleep stages. Dashed lines indicate medians. A mixed-linear model with subject as a random factor revealed slower reaction times in lucid REM sleep. (d) Flowchart detailing the repartition of naps in participants with narcolepsy: the percentage of naps with at least one behavioral response is indicated, and the responsive naps are further divided depending on whether participants reported a lucid dream upon awakening and whether they explicitly recalled responding during the nap. Please note that, after the naps in which participants with narcolepsy responded to stimuli, the majority of participants who were lucid remembered performing the task while those who were not lucid did not. **** : p<0.0001, ***: p<0.001,**: p<0.01, *: p<0.05, all p values are corrected for multiple comparisons using Benjamini–Hochberg procedure.

In order to ensure that participants actually performed the lexical decision task while asleep, we next computed subject-level accuracy scores (Figure 2B). Note that we did not have enough responsive trials per participant to perform this analysis in REM sleep in HP and in N3 sleep in NP. Both HP and NP performed the task significantly more accurately than chance level in all sleep stages, with median accuracy above 71% (HP: Wake 93.7%, *p* < .0001; N1 85.4%, *p* = .0002; N2 83.3%, *p* = .008. NP: Wake 87.9%, *p* < .0001; N1 84.1%, *p* < .0001; N2 71.8%, *p* = .0001; non-lucid REM sleep 73.37%, *p* = .003). We observed a significant main effect of the sleep stages on accuracy in both NP (***χ***²(4) = 38.23, p < .0001) and HP (***χ***²(2) = 13.14, p = .002), indicating a decrease in performance from Wake to deeper sleep stages. Accuracy in Wake, N1 sleep, and N2 sleep was higher in HP than NP (*t* = 2.98, *p* < 0.003).

We then wondered if a behavioral hallmark of lexical decision task during wakefulness, defined by slower response times (RTs) for pseudo-words than for words^21^, could be found in our sleeping participants. For both NP and HP, we found a main effect of both sleep stage (NP: ***χ***²(4) = 82.5, *p* < .0001 ; HP: ***χ***²(2) = 20.7, *p* < .0001) and stimulus type (NP: ***χ***²(1) = 36.9, *p* < .0001; HP: ***χ***²(1) = 59.3, *p* < .0001) on RTs; crucially, there was no significant interaction effect between these two factors (NP: ***χ***²(4) = 7.3, *p* = .1; HP: ***χ***²(2) =2.3, *p* = .32), suggesting that the effect of stimulus type was similar across all sleep stages (Figure S2). Reponses to pseudo-words were on average 130ms slower than responses to words in NP (median: 1.42s), and 120ms slower in HP (median: 1.28s). This was also the case for each sleep stage independently. In NP, responses were faster during wakefulness than during sleep (median RT: Wake, 1.36s *vs* N1 sleep 1.56s, *p* = .034; N2 sleep 1.59s, *p* = .0001; non-lucid REM, 1.49s, *p* = .0001), whereas no significant differences were found between sleep stages (Figure 2C). A similar pattern was observed for HP, including significantly shorter reaction times in Wakefulness and N1 sleep than in N2 sleep (Wake v*s* N2 sleep: *t* = 4.1, *p* < .0001; N1 *vs* N2 sleep: *t* = 2.4, *p* = .025).

We finally assessed whether lucid and non-lucid REM sleep differed on the behavioral level. Only NP reported lucid dreams upon awakening, in 33/134 naps (24.6%). Similar to non-lucid REM sleep, response rates were higher during ON *vs*. OFF periods in lucid REM sleep (52.7% vs 6%, *z* = 18.04, *p* < .0001). Lucid participants also had better performance than chance (65% vs. chance level at 50%, *p* = .001). Lucidity significantly increased the response rate in REM sleep (*z* = 7.97, *p* < .0001) to a level similar to the one observed in N1 sleep (Figure 2A and Table S1). Interestingly, RT were significantly longer during lucid REM sleep than during wakefulness but also than during other sleep stages (median RT: lucid REM sleep, 2.1s vs. N1 sleep, 1.56s, *p* < 0.0001; vs. N2 sleep, 1.59s, *p* = 0.0001; vs. non-lucid REM sleep, 1.49s, *p* = 0.0024) (Figure 2C). Finally, we found a clear association between lucidity and post-nap recall of having performed the task during sleep (Figure 2D): while 75.8% of lucid naps with at least one behavioral response during sleep were associated with a recall of having performed the task, only 15.5% of non-lucid naps with responses were associated with recall (***χ***²(2) = 36.15, *p* < .0001).

As an interim conclusion, our behavioral results demonstrate that sleepers are able to perceive verbal stimuli, make a lexical decision, and perform an adequate motor response while remaining asleep in all sleep stages (except N3 sleep in HP). The fact that participants’ responses were accurate and slower for pseudo-words than for words suggest that stimuli were processed at a high cognitive level. Overall, these results suggest the existence of transient states that allow responsiveness to external information during sleep, whose frequency and duration depend on sleep stage.

### Electrophysiological markers of higher cognitive states predict responsiveness during ordinary sleep

To explore whether responsiveness during sleep could be explained by an ongoing, richer cognitive state prior to stimulation in non-lucid participants (NP and HP), we computed electrophysiological markers known for distinguishing high versus low cognitive states^22,23^. These markers were previously shown to differentiate patients with unresponsive wakefulness syndrome from patients in a minimally conscious state and healthy participants^22,24,25^, as well as wakefulness and REM sleep from N3 sleep^26^. We included five spectral measures (normalized power spectral densities [PSD] in delta, theta, alpha, beta, and gamma frequency bands), one connectivity measure (weighted symbolic mutual information [wSMI] in the theta band), and three complexity measures (the Kolmogorov Complexity [KC], the Permutation Entropy in the theta band [PE θ], and the Sample Entropy [SE]). Crucially, we computed these markers in the 1000ms time window *before* the stimulus presentation; therefore, these markers reflected the “resting-state” brain dynamics of the participants just before the stimulus presentation, and not the evoked activity of the stimulus or the response.

In order to ensure that these markers would provide meaningful information about the cognitive state of our participants, we first assessed how the markers varied in different sleep stages as a sanity check (NP: Figure 3, HP: Figure S3). As expected, complexity measures and high-frequency PSD decreased from wake to N1 sleep, REM sleep, N2 sleep and N3 sleep (in order). Please note that this descending profile mirrored the response rates in those sleep stages. Moreover, delta PSD varied as expected: more delta was observed in N3 sleep compared to N2 sleep, REM sleep, N1 sleep, and wake (in order). Finally, wSMI was higher in wake compared to sleep. These results demonstrated that these markers can reliably distinguish *the sleep/wake stage* of the participants. For the details of the statistical comparisons between the different sleep stages, for each marker and each group, see tables S2 and S3.

**Figure 3.**
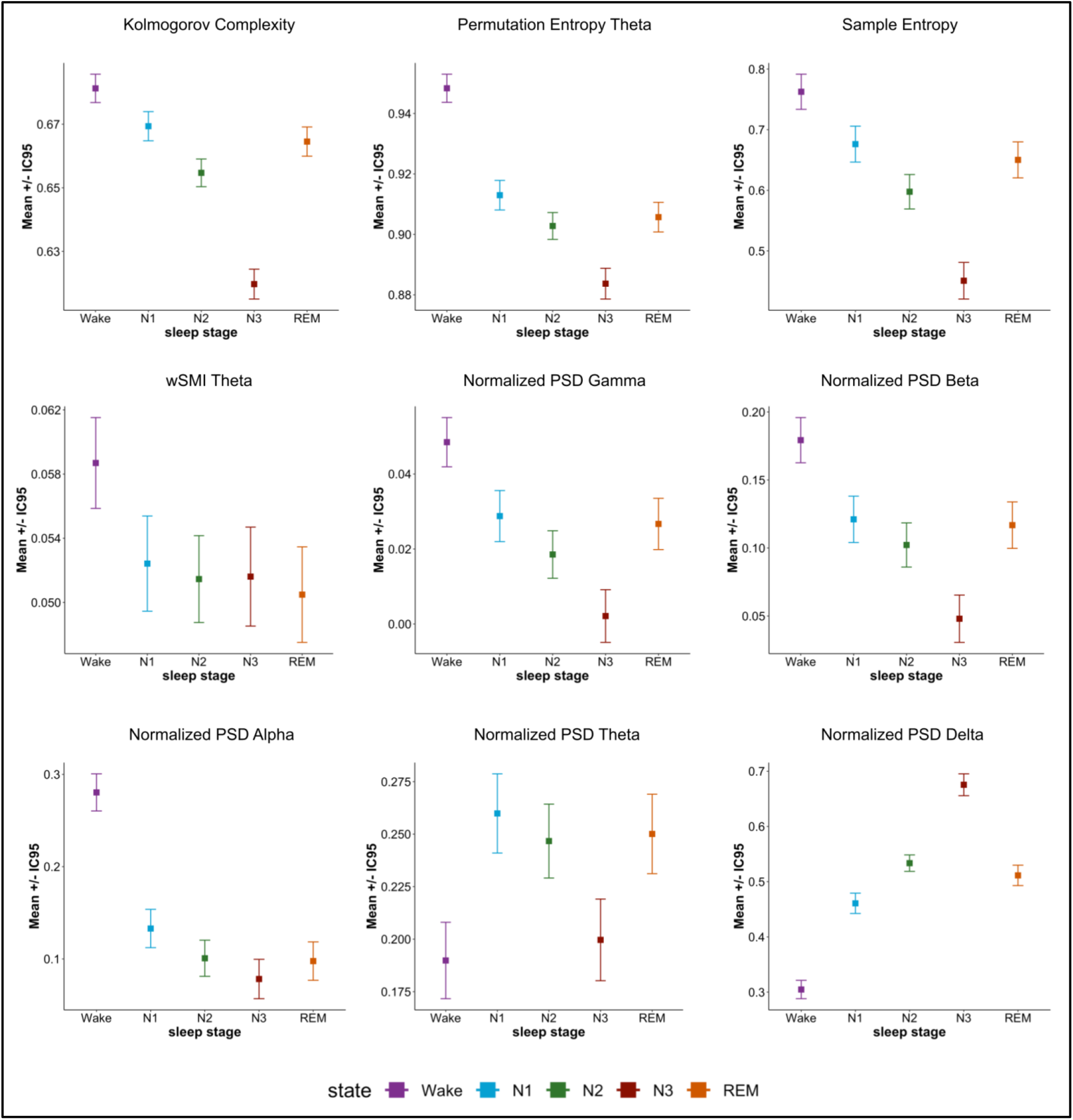
Evolution of electrophysiological markers across sleep stages in participants with narcolepsy. Three complexity measures (the Kolmogorov Complexity -KC, the Permutation Entropy -PE, and the Sample Entropy -SE), one connectivity measure (weighted symbolic mutual information (wSMI) in the theta band), and five spectral measures (normalized power spectral densities (PSD) of delta, theta, alpha, beta and gamma frequency bands) were computed separately for the wake, N1, N2, N3, and REM sleep stages in participants with narcolepsy. The results in healthy participants can be found in Figure S3. Each dot indicates marginal means estimated by a mixed-linear model including sleep stage as an independent variable, EEG marker as the dependent variable, and participant ID as a random variable. Error bars depict 95% confidence intervals. Complexity and high-frequency PSD decreased in sleep compared to wake (wake > N1 > REM sleep > N2 > N3), whereas delta PSD increased with sleep (N3 > N2 > REM sleep > N1 > wake). Details of the statistical comparisons can be found in Table S2.

Next, we assessed how these electrophysiological markers differed in responsive and non-responsive trials (see Table S4 for detailed comparisons). Note that after preprocessing of EEG data, we did not have enough remaining responsive trials in REM sleep for HP and in N3 sleep for NP to conduct this analysis. Figure 4A shows the difference in the estimated marginal means of the z-scored marker values in responsive and non-responsive trials for each sleep stage in non-lucid NP (left panel) and HP (right panel). Positive marker values indicate an increase of the markers in the responsive trials compared to non-responsive trials whereas negative marker values signify a decrease in the responsive trials. Our analysis revealed similar patterns of variations in non-lucid NP and HP, including an increase in the EEG complexity and in the high-frequency PSD, and a decrease in the delta PSD in responsive trials vs non-responsive trials. Connectivity, as assessed by wSMI, did not differ in the two conditions. Importantly, marker values in responsive trials during sleep were never at the same level as wake (even in non-responsive trials), implying that the sleep state was not contaminated by short awakenings (Figure S4). Scalp topographical analyses were not informative given the small number of electrodes (n=10).

**Figure 4.**
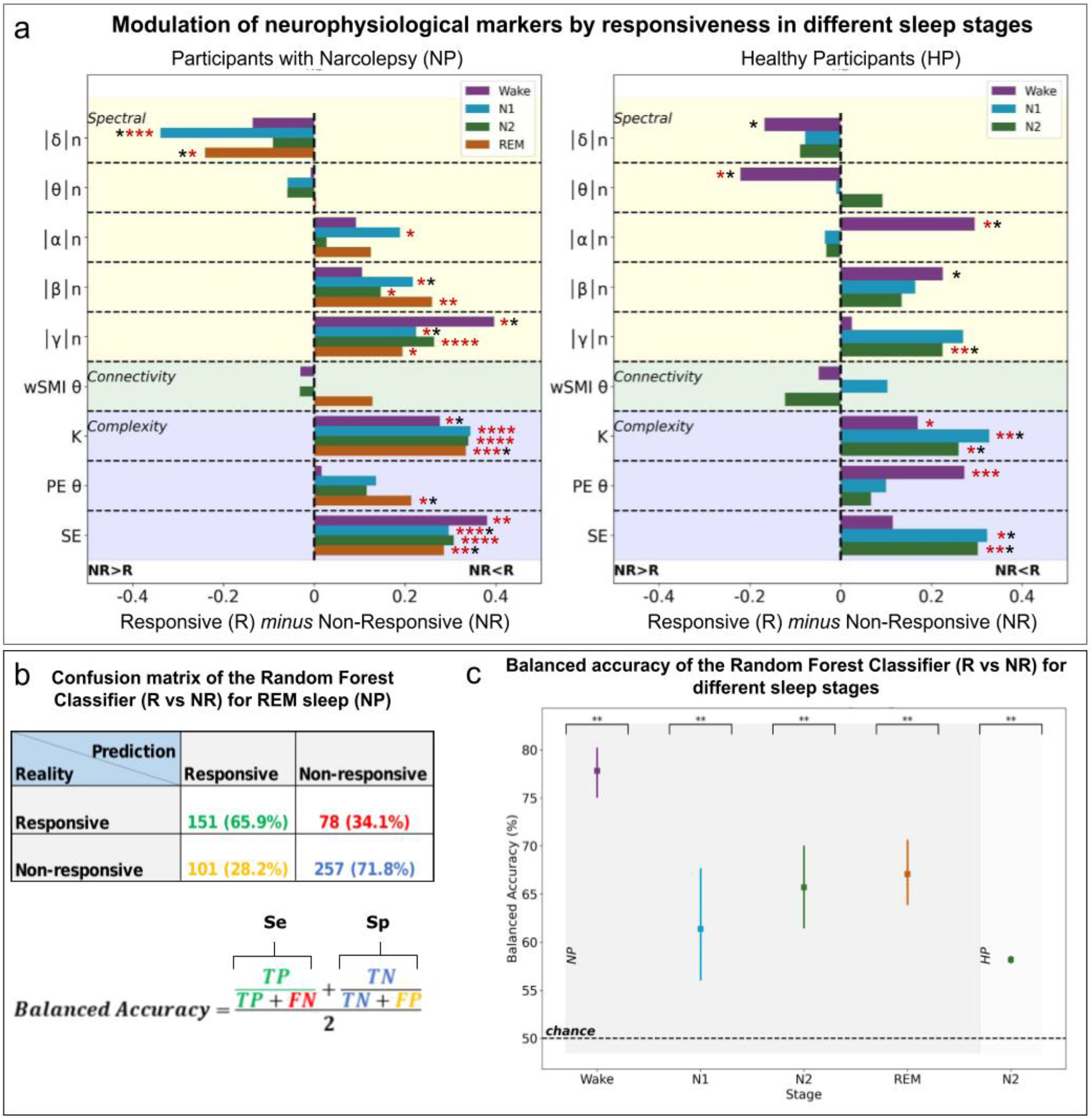
Consciousness markers computed before stimulus presentation predict responsiveness to stimuli in each non-lucid sleep stage. (a) Univariate analysis; After the z-score transform of marker values, we subtracted the marginal estimated mean of non-responsive trials (NR) from responsive (R) trials for each marker and each stage. Almost all markers showed a variation in the direction corresponding to increased conscious processing when contrasting responsive trials to non-responsive trials (e.g.: increased EEG complexity and decreased EEG delta power), both in participants with (left) and without narcolepsy (right). Note the similarity with Figure 3 in Sitt et al., (2014) which contrasted conscious to non-conscious states in patients suffering from disorders of consciousness. (b) and (c) Multivariate analysis. We fed a random forest classifier with these 9 EEG markers and trained it to classify R trials versus NR ones using a 10-fold cross-validation method. A confusion matrix for REM sleep stage in non-lucid naps of participants with narcolepsy is shown in (b), with a description of the balanced accuracy measure that we computed to take unbalanced datasets into account. The confusion matrix for each stage and group can be found in Table S5. TP: True Positives (responsive trials classified as responsive). TN: True Negatives (non-responsive trials classified as non-responsive). FP: False Positives (NR trials classified as responsive). FN: False Negatives (R trials classified as non-responsive). Balanced accuracy scores are plotted in (c) for different sleep stages, both for participants with narcolepsy (Wake, N1, N2, REM sleep; left) and without narcolepsy (N2, right), with the corresponding statistical significance against chance-level (approximated by 500 permutations). Note that: (i) responsiveness to stimuli could be predicted for each sleep/wake stage in participants with narcolepsy (NP); (ii) the classifier trained with data from participants with narcolepsy could generalize to healthy participants (HP), as shown by significant decoding of responsiveness in N2. **** : p<0.0001, ***: p<0.001,**: p<0.01, *: p<0.05, red stars indicate significance after FDR correction for 72 comparisons.

To further explore the predictive power of these EEG markers on responsiveness, we trained a random forest classifier using a multivariate combination of the markers collected in non-lucid NP and did so independently for each sleep stage. We then tested whether this classifier could predict responsiveness on a trial-by-trial basis in both NP (using a classical stratified cross-validation procedure) and HP trials (in N2 sleep). The balanced accuracy score was above 60% for all sleep stages in NP non-lucid naps (reaching 67% for REM sleep) and reached 58% for N2 sleep in HP (Figure 4C). All balanced accuracy scores were significantly different than the chance level computed by a 500-permutation procedure *(p* < .002 for all stages in NP, and *p* = .006 for N2 sleep in HP), with a mean balanced accuracy score of permutation trials very close to 50% for all stages (Table S5).

In sum, the EEG results suggested that a particular brain state prior to the stimulation, characterized by increased complexity and faster oscillations, allowed responsiveness during sleep. A multivariate combination of these markers predicted the presence/absence of response in a trial-by-trial level. The fact that the markers varied with responsiveness similarly in non-lucid NP and HP and that the classifier trained with NP data could classify responsive trials in HP better than chance strongly suggest that the same brain dynamics underlie responsiveness in both participants with and without narcolepsy (in non-lucid sleep).

### Lucid dreaming as a proxy for inferring conscious processing of responded stimuli in other sleep stages

To further investigate the specificities of lucid REM sleep in NP, we first compared the electrophysiological markers between responsive and non-responsive trials in this condition. Interestingly, none of these markers differentiated responsive from non-responsive trials in lucid REM sleep (all uncorrected *p* > 0.05) (Figure 5A and Supplementary Table 6). To check that this null result was not due to a lack of statistical power in our frequentist approach, we conducted, for each marker independently, a supplementary Bayesian analysis (see Table S6). We computed the Bayes Factor (BF) of our full mixed linear model and compared it to the one of a “null-model” with only the random effect. All BF were inferior to 1 (ranging from 0.21 to 0.08). Such values of BF indicate moderate (<0.33) to strong (<0.1) evidence for the null model^27^, suggesting a true absence of difference between responsive and non-responsive trials in lucid REM sleep.

**Figure 5.**
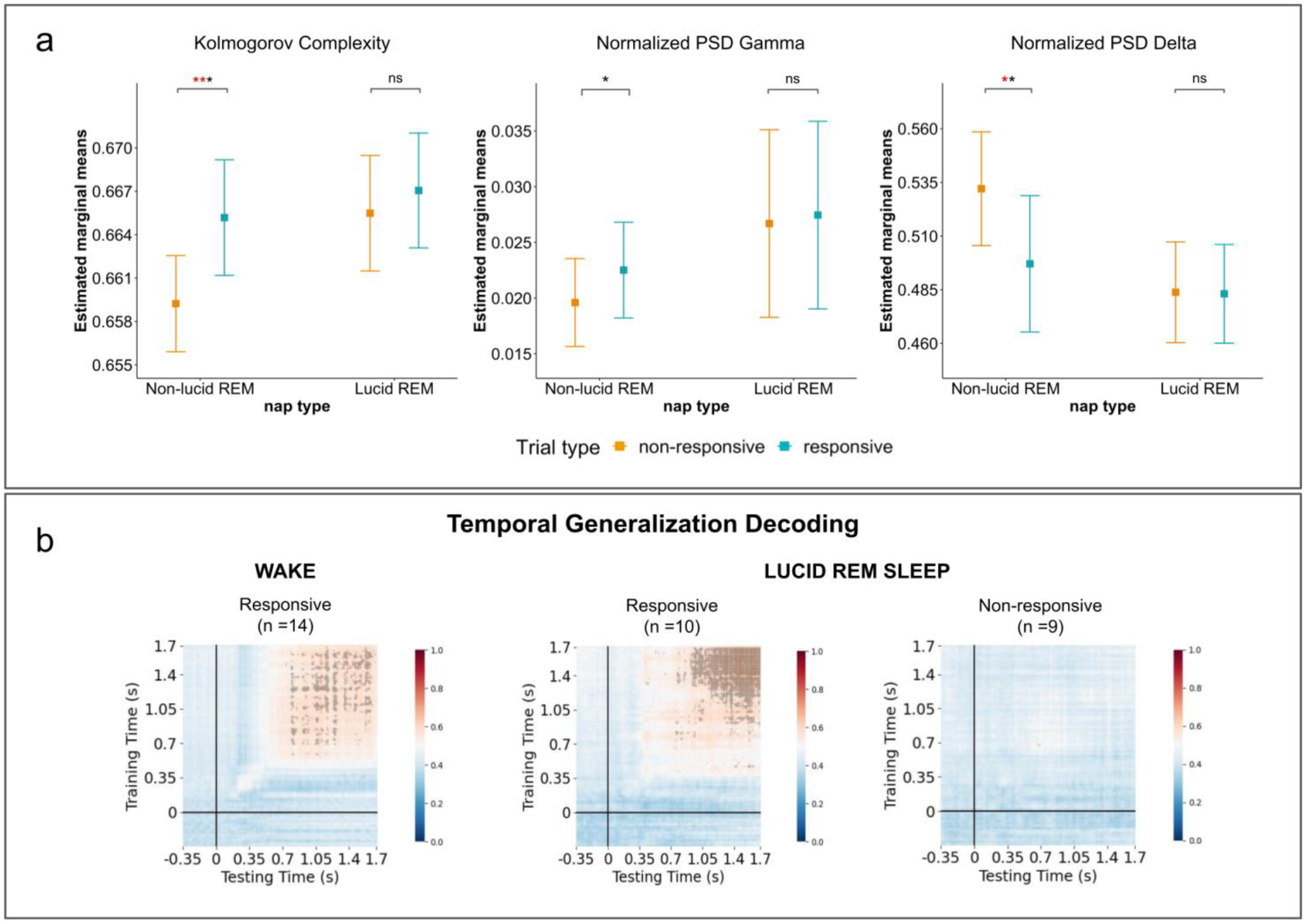
Effect of lucidity on EEG markers and response to stimuli in participants with narcolepsy. (a) The top panel shows Kolmogorov complexity (left), normalized gamma PSD - norm-*gamma* (middle), and normalized delta PSD *-norm-delta* (right) prior to stimuli onset as a function of whether the stimulus will be followed by a behavioral response (in blue) or not (in orange), for lucid and non-lucid REM sleep in participants with narcolepsy. Kolmogorov complexity and norm-gamma were significantly higher for responsive trials compared to non-responsive trials in non-lucid naps for all participants. Conversely, the norm-delta was significantly lower in responsive trials in non-lucid naps. No such differences were observed in lucid naps, suggesting a ceiling-effect for cognitive richness markers in lucid naps. Overall, Kolmogorov complexity and norm-gamma were higher, and norm-delta was lower in lucid naps compared to non-lucid naps irrespectively of the responsiveness. Time-generalization decoding of stimulus-related brain activity compared to baseline brain activity, in trials with (top) and without (bottom) response, in Wake (left) and Lucid REM sleep (right). The logistic regression classifier was trained on each time point and then tested on all the time points to obtain a generalization pattern. Each intersection point of a training time and a testing time shows the AUC (Area under the receiver operator curve) of the classifier. Time points with an AUC>0.5 and that are statistically significant are outlined in black (two-sided non-parametric sign test across subjects with FDR correction for 41 616 comparisons, *p* < 0.05). Note that in trials with a behavioral response, we observed for both Wake and Lucid REM sleep short diagonal pattern suggestive of a ballistic transient chain of distinct processing stages, followed by a squared-shape pattern revealing a late, stable, and sustained stage of processing which has been previously associated with conscious access. **** : p<0.0001, ***: p<0.001,**: p<0.01, *: p<0.05, ns: non-significant, red stars indicate significance after FDR correction for 15 comparisons.

We next investigated how the marker values in lucid REM sleep differed from the ones in non-lucid (ordinary) REM sleep. We observed higher complexity (sample entropy) and normalized PSD of gamma, as well as lower normalized PSD of delta values in lucid trials compared to non-lucid trials. Interestingly, statistical analyses (both frequentist and Bayesian) restricted to only responsive trials revealed similar values in lucid and non-lucid conditions for all markers, indicating comparable brain activity during responsive trials between non-lucid and lucid REM sleep (Figure 5A and Table S7).

In sum, lucid REM sleep was characterized by a systematic increase in EEG markers of higher cognitive states, irrespective of behavioral responsiveness to the task, with a pattern of markers similar to the one observed in non-lucid/responsive trials (i.e. faster oscillations and higher complexity compared to non-lucid/non-responsive REM trials). This is evocative of a ceiling effect for marker values in lucid REM sleep, indicating a steadily high-cognitive state during this condition.

This neurophysiological profile combined with the subjective report of having performed the task during sleep (see behavioral results) suggest that NP *consciously* processed the stimuli *when* in lucid sleep. Several signatures of conscious processing have been described in the literature, such as the late P3b component in evoked related potentials^28–30^ or the square-like shape pattern in the temporal generalization method^31,32^. As the latter was more appropriate to the specificities of our dataset (e.g., unbalanced trials), we explored whether the pattern of temporal generalization supports our hypothesis of a conscious processing of external stimuli during lucid REM sleep. In brief, this analysis tests how stimulus-induced brain activity differs from baseline activity; it consists in training a linear classifier at each time-point to differentiate stimulus-present versus stimulus-absent epochs and testing its performance for all the other time-points (for example, training the classifier at t=2 and testing its ability to correctly classify at t=1,2,3,4,5,…, obtaining thus a whole matrix of performance for each training time point/testing time point). We found that responsive trials during lucid REM sleep were associated with the expected square-like shape pattern starting from 350ms post-stimulus presentation. Such a pattern reflects a late, stable, and sustained processing stage that has been previously related to conscious access^31–33^ (Figure 5B). Importantly, this pattern was very similar to the one observed in responsive Wake trials, indirectly supporting our hypothesis that NP are conscious of the stimuli presented during responsive trials in lucid REM sleep. In contrast, we did not find any discernible decoding pattern for non-responsive trials in lucid REM sleep, suggesting that NP are not conscious of external stimuli when they do not respond. This result might seem at odds with our previous observation that marker values computed *prior* to the stimulation were similarly high in responsive and non-responsive trials (Figure 5A) in lucid NP. It suggests that high marker values are indicative of a rich cognitive state, which is permissive (but not necessarily sufficient) for responsiveness during sleep. We hypothesize that missed trials (no responses) during lucid REM sleep are due to a competition for attentional resources between external stimuli and internal percepts (higher attention to the ongoing dream)^1^, resulting in a dual task. This hypothesis is consistent with the longer reaction times (a typical hallmark of dual task) observed in lucid REM sleep (see behavioral results).

Lucid REM sleep trials were therefore associated with: (I) a subjective report of having performed the task while sleeping; (II) a systematic increase in EEG markers of higher cognitive states; (III) an electrophysiological signature of conscious processing of external stimuli in responsive trials (temporal generalization pattern), and (IV) longer reaction times, suggesting that participants were engaged in a dual task during which external information (outside world, including verbal stimuli) and internal information (ongoing dream) compete for attention. All of these findings not only hint that lucid participants could integrate and respond to external stimuli during sleep, but also that this process was conscious.

## DISCUSSION

Our results provide compelling evidence that sleeping humans present, in all sleep stages, transient windows of sensory connection with the outside world during which they process external information at a high-cognitive level and are able to exhibit a behavioral response. Until now, behavioral responsiveness had only been demonstrated during some light sub-stages of N1 sleep (sleep onset)^4,15,16^ or in some unique individuals during lucid REM sleep^14^. Our findings go further by demonstrating the possibility for behavioral responsiveness to external stimuli across all sleep stages in ordinary sleep in a large group of participants. Furthermore, we show that these transient windows of sensory connection are associated with specific brain dynamics (faster oscillatory activity and higher signal complexity), which predict responsiveness on a trial-by-trial basis. Finally, for the particular case of lucid REM sleep, we provide strong arguments in favor of a *conscious* processing of *external* information, including the presence of a neural signature of conscious access^31^ in responsive trials and explicit recall of having performed the task during sleep. This finding reveals that lucid dreamers’ enhanced consciousness of their internal world also extends to the external world during REM sleep.

Before further discussing our results, it is important to stress several limitations of our study. First, behavioral responses were assessed by a visual inspection of the corrugator and zygomatic muscle activity on EMG channels. We chose this strategy rather than an automated scoring procedure because the amplitude of muscular responses was highly variable between participants and between trials (even during Wake). However, because behavioral responses were scored while blind to the stimulation period (ON or OFF), the participant identity and the sleeping stage (all channels other than corrugator and zygomatic EMG channels were removed before scoring), any subjective bias during the scoring procedure affected every sleep stage and stimulation period the same way. Second, the fact that this experiment was done over the course of several daytime naps limited the quantity of N3 sleep that we could record, preventing us from assessing the presence of behavioral responses during N3 sleep in healthy participants or to measure our neurophysiological markers in responsive N3 sleep trials in participants with narcolepsy. Further studies are necessary to better assess behavioral responsiveness during N3 sleep, and more generally during night-time sleep (instead of daytime naps). Third, the whole-brain connectivity as assessed with wSMI at theta frequency did not differ between responsive and non-responsive trials in any of the sleep stages. A recent study^15^ showed that whole-brain connectivity might not be as relevant as fronto-temporal connectivity when predicting responses in N1 sleep. Our small number of electrodes limits the interpretation of this connectivity metric. Recording with more electrodes in the frontal and temporal regions are needed to further investigate the link between the functional connectivity and responsiveness in sleep. Fourth, we used post-nap subjective reports to determine lucidity instead of the gold-standard, objective signal of lucidity^18,34,35^. Nevertheless, we also collected an objective lucidity signal (successive corrugator and zygomatic contractions) that substantially matched participants’ subjective reports upon awakening (supplementary results), confirming the reliability of subjective reports in determining participants’ lucidity. Additionally, we only obtained lucid naps in patients with narcolepsy. Therefore, our results for lucid REM sleep need confirmation in lucid healthy participants.

Although the response rate was minimal during OFF periods (compared to ON periods), it was still greater than zero, which might appear surprising in the absence of stimuli. This can be due to several factors: (i) participants might have had spontaneous contractions, (ii) they might have dreamt about the task and contracted their muscles in response to a dreamt auditory stimulation, or (iii) we might have over-estimated the contraction rates. Spontaneous single contractions called ‘twitches’’ are common during REM sleep. However, we only considered two or more successive contractions as responses, eliminating all twitches. Moreover, as mentioned before, behavioral responses were assessed while blind to the sleep stage and to the stimulation period (ON vs OFF). Even if blind scorers had false alarms when detecting contractions, this bias should be uniformly distributed in all sleep stages and stimulation periods. Therefore, any differences in the response rates between ON and OFF periods reflect a genuine effect.

One might argue that the behavioral responses we observed during sleep occurred during brief episodes of wakefulness. Yet, all trials containing a micro-arousal (before and/or after the stimulation) were excluded from all analyses to ensure that participants were indeed asleep while responding, at least according to the well-accepted sleep scoring rules^20^. However, recent studies showed that local-sleep phenomena can be observed during wake, suggesting that the discrete frontiers between wake and sleep might be fuzzier than the international sleep criteria would suggest^36,37^. In the same way, it is possible that participants had ‘local wake events’ allowing them to respond to external stimuli while sleeping. Our current gold-standard sleep scoring guidelines are not suited to detect such subtle variations in brain dynamics. In this sense, it is important to stress that some of the neurophysiological markers used in this study could also be interpreted as markers of arousal (for example, more power in fast frequencies and less delta power). Our study could thus precipitate the development of finer-grained sleep scoring which better captures cognitive capacities including behavioral responsiveness in the wake-sleep continuum.

One interesting yet expected finding was that participants with narcolepsy had higher contraction rates compared to participants without narcolepsy. Given their tendency to fall asleep in unconventional situations, they might have acquired the capacity to remain connected with their surroundings while sleeping. Alternatively, a reduced muscle atonia compared to healthy controls^38^ might enable participants with narcolepsy to respond more during sleep. Finally, patients with narcolepsy have many symptoms reflecting a sleep-wake instability (e.g., sleep paralysis, hypnagogic hallucinations), and might thus be more prone to experience “local wake events”, enabling them to respond more frequently to external stimuli during sleep.

Our results enhance our understanding of the lucid dream phenomenon and of its neural correlates^35,39^. We found modifications in spectral power (increase in normalized PSD of gamma and decrease in normalized PSD of delta) as well as an increase in signal complexity (Sample Entropy) during lucid REM sleep, compared to ordinary (non-lucid) REM sleep. Interestingly, a reduction in low-frequency spectral power coupled with an increase in signal complexity were also found in a recent study^40^. Our low-density EEG montage (10 electrodes) did not allow for a more precise topographical description of these modifications, but this could be clarified in future studies. More importantly, we provide strong evidence that, in the case of lucid participants, the stimuli in our task were perceived in a conscious manner. Indeed, participants reported having heard the stimuli and having performed the task upon awakening. On top of this subjective report, which is the gold standard to assess conscious access, they also exhibited stable and sustained brain activity in response to stimuli. Such neural responses have been previously shown to reflect conscious perception^31–33^. Finally, neural metrics computed prior to the stimulation indicated a richer neural state presumably allowing responsiveness. Evidence from literature demonstrates that these rich neural states differentiate conscious from unconscious humans (for example, patients with unresponsive wakefulness syndrome^22,24,25^). Taken together, these observations strongly imply that lucid participants consciously responded to stimuli while asleep. These results extend our understanding of lucid dreaming at the cognitive level, by showing that lucid dreaming is not only characterized by a reemergence of metacognitive and volitional capacities^41,42^, but also by a capacity to consciously process external information.

To what extent were non-lucid sleepers conscious when responding to stimuli? The answer is not as clear as in lucid dreamers since non-lucid dreamers typically could not recall having performed the task during sleep. Moreover, due to the insufficient number of trials, we could not perform temporal generalization decoding in these participants and therefore could not investigate neural responses to stimuli. However, there are several lines of evidence that favor conscious processing in this population. First, neurophysiological markers computed before the stimulation in responsive trials were similar to the ones in lucid participants. This suggests that their neural state in the responsive trials was comparable to the one in lucid participants. Furthermore, the unconventionality of the response modality (corrugator or zygomatic muscle contractions) makes the automatization of the task difficult. Indeed, we rarely use these muscles in everyday life to convey binary responses. Finally, reaction times to stimuli exceeded several seconds, a duration much longer than the one (typically around 200ms) classically observed for automatic and unconscious processing^43^. One may wonder why participants would fail to report having done the task if they had consciously performed it. One possibility is that they simply forgot or did not encode this information. Some cases of lucidity amnesia have already been described in the literature^34,44^. Moreover, some of our participants signaled their lucidity using mixed contractions and did not remember doing so nor being lucid at awakening. We could also hypothesize that the rich neural states presumably allowing responsiveness need to be sustained over a certain period of time in order to be encoded. These rich states might have been less stable in non-lucid participants (as suggested by the difference in neurophysiological markers between responsive and non-responsive trials in non-lucid participants (not found in lucid ones), preventing episodic memory encoding and thus subjective reports.

The standard, binary view of sleep/wake states assumes that we would be either awake or asleep. Overall, our findings suggest that this view does not account for the richness and high variability within each of these states. Our results show that access to external information is a fluctuating phenomenon that might vary even in traditionally defined states of consciousness (e.g., a given sleep stage). We could imagine sleep and wake as a continuum of stages whose physiology is more (e.g. wake) or less (e,g, N3 sleep) favorable for the emergence of the rich neural states that enable conscious access and behavioral response to external stimuli^46^. In this sense, it is interesting to note that the values of the neurophysiological markers computed at the sleep stage level mirrored the response rates at these given sleep stages.

Our study opens the way for many exciting studies investigating sleepers’ cognitive capacities and their associated phenomenology. With some small modifications of the current approach (e.g. implementing a second probe about a participant’s current mental state), we could assess metacognition during responsive moments (e.g., do sleepers know that stimuli come from the outside, or do they integrate them in their dream?). By tracking how the neurophysiological markers indexing a rich cognitive state fluctuate in real time and by sending stimuli depending on their values, we could test the causal relationship between the neural state and responsiveness. Moreover, as the brief windows of reactivity in sleep can be predicted from EEG signal, they could be targeted to test the possibility for real-time communication with sleepers, not only in some unique individuals who experience lucid dreams^14^, but in all sleepers across all sleep stages. Such two-way communication with sleepers may open novel research avenues, allowing inquiries about sleepers’ mental states in different sleep stages. This new method could also have clinical applications, for example in the medical treatment of patients suffering from post-traumatic stress disorder, by communicating with them during recurrent nightmares as a way to relieve them, or for bringing mechanistic insights into the puzzling mismatch between subjective wake perception and objective sleep markers in patients suffering from paradoxical insomnia. By demonstrating the existence of windows of behavioral responsiveness across all sleep stages, our study provides a new tool for unlocking the mystery of what happens in sleepers’ minds.

## MATERIAL AND METHODS

### Participants

#### Participants with narcolepsy

Thirty participants with narcolepsy were recruited for this study (14 women, mean age: 35 ± 11 years) from the patients followed in the National Reference Center for Narcolepsy in Pitié-Salpêtrière University Hospital. Twenty-four of them (80%) were frequent lucid dreamers who reported more than 3 lucid dreams per week on average (others reported less than 1 lucid dream per year). Participants met the international criteria for narcolepsy^46^, including (i) excessive daytime sleepiness occurring daily for at least 3 months; (ii) a mean sleep latency lower than or equal to 8 min and two or more sleep onset REM sleep periods on the multiple sleep latency tests (5 tests performed at 08:00, 10:00, 12:00, 14:00, and 16:00; and (iii) no other better cause for these findings, including sleep apnea syndrome, insufficient sleep, delayed sleep phase disorder, depression, and the effect of medication or substances or their withdrawal. They were required to pause their medication for the day of the experiment to facilitate sleep onset. We recruited patients with narcolepsy type 1 (n= 17, with clear cataplexy or hypocretin deficiency) and type 2 (n= 13, no cataplexy or hypocretin deficiency). Among the 30 participants, 3 (2 women) were discarded from the analyses because of technical issues affecting the recordings. In total, data from 27 participants with narcolepsy (21 frequent lucid dreamers) were analyzed in this study.

#### Healthy participants

Twenty-two healthy participants (all non-lucid dreamers) were recruited for this study (10 women, mean age: 24 ± 4 years). They had no or little experience with lucid dreaming (less than two lucid dreams in their lives). They had no sleep disorder and were in good shape, as assessed by a sleep clinician. To further facilitate sleep onset, we asked participants to sleep about 30% less than usual during the night preceding the experiment (either by going to bed later or waking up earlier) and to avoid stimulants on the day of the experiment. Fourteen went through the experiment in the morning and eight of them went through the experiment in the afternoon. One participant was discarded from the analysis because of technical issues affecting the recordings.

All participants were native French speakers and gave written consent to participate in the study. The protocol had been approved by the local ethics committee (CPP Ile-de-France 8). Participants with and without narcolepsy were paid €200 and €70 respectively, as compensation for their participation in the study (participants with narcolepsy also took part to an unrelated experiment the following day; the results of this second study are not described here).

### Experimental design

In this study, we tested participants’ ability to perceive, discriminate, and respond to auditory stimuli while asleep. Participants lied in a bed in a sound attenuated room in the sleep unit. They were asked to perform a lexical decision task in which words and pseudo-words were verbally presented in a pseudo-randomized fashion. Participants with narcolepsy went through five 20-min naps, with an 80-min break between each nap (Figure 1A). Before the experiment, participants underwent a short training (10 min) to familiarize themselves with the type of stimuli and the task (10 repetitions). Each nap session contained ten “ON” stimulation periods during which 6 stimuli (3 words and 3 pseudo-words) were presented every 9 to 11 seconds on top of continuous white noise. Each stimulus was presented only once in the entire experiment. The “ON’’ stimulation periods were separated by 1 min non-stimulation periods (OFF periods) during which only white noise was presented. Following a previously validated response paradigm during sleep^14^, participants were instructed to decide whether the stimulus was a word or a pseudo-word and indicate their response by making three, brief, successive contractions of either the corrugator (frowning) or the zygomatic (smiling) muscles, depending on the stimulus type (e.g., contracting the corrugator if they heard a pseudo-word and the zygomatic if they heard a word). The muscle-stimulus association was counterbalanced across participants. Importantly, the stimulation started when the subjects were still awake, but participants were explicitly authorized to fall asleep while performing the task. They were asked to perform the task before falling asleep, if they woke up during a nap, and if they heard the stimuli in their sleep. If participants were lucid dreaming but did not hear any stimuli (word or pseudo-words), they were instructed to communicate their lucidity with a “mixed” signal, alternating a single corrugator muscle and a single zygomatic muscle contraction. Note that we chose not to use the gold-standard method to signal lucidity here (Left-Right-Left-Right ocular code) for three reasons: i) the ocular code ‘pollutes’ the EOG channel, which might lead to bias when scoring REM sleep, ii) several lucid dreamers with narcolepsy explicitly told us that facial codes were easier to perform, less disturbing of the ongoing dream, and less awakening than the ocular code, and iii) our experiment required three different codes (one for each stimulus type and one for signaling lucidity if no sounds were heard). After each nap, participants were awakened by an alarm that rang until they pressed a button. They were asked to report ‘what was going through their mind’ before the alarm and indicate whether i) they had a lucid dream, ii) they communicated their lucidity with the mixed-signal, iii) they heard the stimuli during the nap, iv) they responded to the stimuli, and v) they remember any stimuli (word or pseudo-word) from the nap (free recall). Finally, participants performed an old-new recognition task, during which they were presented with stimuli they heard during the preceding nap and new stimuli that were never presented during the experiment. Participants had to indicate whether they had heard the stimuli during the preceding session with one of the following responses: 1: I heard it from the dream (for example, a person from their dream saying the word), 2: I heard it from the outside world (pronounced by the computer), 3: I am not sure I heard it, 4: I am sure I did not hear it. They responded by pressing the corresponding button without any time pressure. The four options were explained to the participants during training, prior to the first session.

Healthy participants went through the same procedure except that the 5 naps were combined into a single, longer, 100-min daytime nap.

### Stimuli

Stimuli were French words and pseudo-words pronounced by a female voice, taken from the MEGALEX database^45^. All stimuli were controlled for their duration (690ms) and the words were controlled for their frequency and valence. Five distinct lists (one for each nap session) of sixty stimuli (thirty words and thirty pseudo-words) were created for each participant in a randomized fashion. Participants heard each stimulus only once during the day. Stimuli were presented through speakers using the Psychtoolbox extension^47^ for MATLAB (The MathWorks). Stimuli were played every 9–11 s (random uniform jitter) after a 60 second OFF period (without stimuli). Button-press responses in the old-new recognition task were collected through a regular keypad.

### Electrophysiological recording

Electroencephalography (EEG, 10 channels: Fp1, Fp2, Cz, C3, C4, Pz, P3, P4, O1, O2, referenced to the A2; 10–20 montage), electrooculography (EOG, 2 channels, positioned above the right superior canthus and the left inferior canthus), electromyography (EMG, 1 channel on chin muscle for sleep staging, 1 channel on zygomatic and 1 channel on corrugator muscles for recording participants’ behavioral responses) and electrocardiography (EKG, 1 channel) were continuously recorded during the nap sessions. All signals were recorded simultaneously at a 2048 Hz sampling rate. EEG data were amplified through a Grael 4K PSG:EEG amplifier (Medical Data Technology, Compumedics Ltd, Australia).

### Sleep scoring and identification of muscular responses

#### Sleep scoring

Sleep stages were scored offline by a certified sleep expert according to established guidelines^49^ using Profusion software (COMPUMEDICS, Medical Data Technology). For scoring, the EEG and EOG signals were filtered between 0,3 Hz and 15 Hz, the EMG and EKG signals were filtered between 10Hz-100Hz and 0,3Hz-70Hz respectively. A 50 Hz notch filter was applied on all channels. Sleep scoring was visually performed on 30-second time epochs, each scored as wakefulness, N1, N2, N3, or REM sleep, according to the AASM international rules. Micro-arousals were scored when alpha rhythm was present during more than 3 sec and less than 15 sec (if longer, the epoch was scored as wake) and, in REM sleep, when there was an increase in chin muscle tone in addition to the alpha rhythm. Trials containing micro-arousals were excluded from further analyses. A nap was considered lucid based on the subjective report (if the participant reported having a lucid dream during the nap). In this case, all REM sleep epochs of this nap were then considered as lucid REM sleep. Note that healthy participants never reported having a lucid dream.

#### Identification of muscular responses

The recording of the nap was divided into 120 mini-epochs of 10 seconds. The sleep stage for each mini-epoch was defined by the sleep score of the corresponding 30-second epoch. Mini-epochs containing a micro-arousal were discarded from the analyses. The presence of zygomatic or corrugator muscle contractions was assessed visually, looking offline at the EMG signal for each mini-epoch. Importantly, the scorer was blind to the sleep stage and to whether a stimulus was presented during the mini-epoch (corresponding to an ON period) or not (corresponding to an OFF period). Muscle contractions were considered as a response if they contained at least two consecutive contractions. Single contractions were considered as a twitch and scored as a no-response. To ensure the quality of the scoring, 10% of the data was later re-evaluated by another blind scorer who showed 84% consistency with the first scorer.

### EEG preprocessing and analysis

Only the EEG segments corresponding to the “ON periods” were analyzed.

#### Preprocessing

Raw files were set to a mastoid reference (A2 electrode). We applied two different preprocessing procedures:

##### For calculation of electroencephalographic markers of consciousness and related machine learning classification

Following previous work^22^, raw EEG files were band-pass filtered between 0.5 and 45Hz, with 50Hz and 100 Hz notch filters. Data was down-sampled to 250Hz. Trials were then segmented from - 1000ms to the onset of the stimuli (words and pseudo-words during ON-periods).

The obtained epochs were cleaned, based on their voltage maximum peak-to-peak amplitude, using a fully automatic procedure with the *autoreject*^48^ algorithm. The *Python*^49^ implementation of the *autoreject* algorithm allows for the automatic calculation of an optimal global rejection threshold for a set of epochs, using a cross-validated machine learning algorithm. For each wake/sleep stage in our data (Wake, N1, N2, N3, and REM sleep), we calculated a separate global rejection threshold (the same for all participants in each group for a given sleep/wake stage) and we rejected all trials with at least one EEG channel exceeding the given threshold. Note that this drastic rejection method was associated with high rejection rates but ensured the quality of our data. More conservative automatic cleaning methods such as interpolation of bad channels were not applicable to our 10 channels EEG montage.

##### For Temporal Generalization Decoding against baseline analysis

Raw EEG files were band-pass filtered between 0.1 and 20Hz, with 50Hz and 100 Hz notch filters. Data was down-sampled to 250Hz. Trials were then segmented from -350ms to +1700ms relative to the onset of the stimuli (words and pseudo-words during ON-periods). The obtained epochs were then cleaned using the same automatic procedure described above.

All trials were labeled as belonging to a particular *sleep/wake stage* (Wake, N1, N2, N3, or REM sleep) according to the sleep scoring described above (corresponding 10s mini-epoch), as being *responsive* or *non-responsive* according to the presence or absence of a valid behavioral response (corrugator or zygomatic muscle contraction), and as *lucid* or *non-lucid* according to the global label of the nap (cf. above).

#### Calculation of electroencephalographic markers tracking consciousness modifications

Previous work has shown that consciousness state modifications can be tracked using different spectral, connectivity, or complexity measures derived from the scalp or intracranial electroencephalographic recordings. By combining these markers, it is possible to distinguish conscious participants, patients in a *minimal consciousness state*, and patients with unresponsive wakefulness syndrome^22,25^. These measures can also differentiate sleep stages (REM sleep and wakefulness *versus* N3)^26^ and track consciousness modifications related to psychedelics or meditation^23^.

In our study, we selected 3 types of measures among those markers:

- <u>Spectral measures:</u> we computed the normalized power spectral densities (PSD) in delta (1-4 Hz), theta (4-8 Hz), alpha (8-12 Hz), beta (12-30 Hz), and gamma (30-45) frequency bands.
- <u>Connectivity measures:</u> we computed the weighted symbolic mutual information (wSMI), a functional connectivity measure capturing linear and non-linear coupling between sensors, which relies on the symbolic transformation of the EEG signal. We computed the wSMI in the theta band (4-8Hz)^24^. The choice of the theta frequency band was based on previously reported results^22,24^, showing that the wSMI calculated on this frequency band was the most efficient in detecting residual consciousness in brain-injured patients with a *disorder of consciousness*.
- <u>Complexity measures:</u> we computed three different complexity measures, the Kolmogorov Complexity (KC), the Permutation Entropy in the theta frequency (PE θ), and the Sample Entropy (SE).

See Supplementary material of Sitt et. al (2014)^22^ for a detailed description of each measure and its computation. Details regarding the sample entropy can be found in Richman & Moorman (2000)^50^.

Each one of the previously described markers was computed during the 1000ms time window *preceding* the presentation of the stimulus (word or pseudo-word), during the ON-periods, independently for each subject, trial and for every electrode (n = 10) or pair of electrodes (n = 45) for the wSMI. A wSMI global score for each electrode was computed by calculating the median connectivity of each electrode with all the other electrodes. Finally, for each subject and each trial, each marker was summarized by calculating the mean across channels, resulting in a single scalar per marker per trial.

### Prediction of responsiveness using a decision tree algorithm

We aimed at predicting, independently for each sleep/wake stage, if a given trial would contain a response or not based on the consciousness markers computed during the 1000ms time period preceding the stimulus presentation. We used a Random Forest algorithm, a classification algorithm consisting of many decision trees. This algorithm implements bootstrapping and feature randomness when building each tree, which ensures the construction of an uncorrelated forest of trees. Since the different trees in the forest are uncorrelated, their global prediction by committee is more accurate than that of any individual tree. Random Forest has shown to be among the best currently used machine learning classifiers, in a very wide range of different datasets (n=112) from several research fields^51^, outperforming other choices as SVM classifiers.

We conducted an independent analysis for each sleep/wake stage. For each trial, the classifier was provided with 10 features, as well as the label (“responsive” versus “non-responsive”) of the trial. The 10 features were the 9 consciousness markers described in the previous section and the subject identity. The Random Forest classifier was composed of 100 estimators (trees). Since our data was unbalanced in terms of the number of responsive trials compared to non-responsive ones, the weights of each class were adjusted in an inversely proportional manner to class frequencies.

Two different training/testing strategies were used:

- For the participants with narcolepsy, we used for each stage a standard 10-fold stratified cross-validation procedure. In each fold, data was split into training (9/10 of the trials) and testing (1/10 of the trials) sets, in a manner that preserved class frequencies in each split. Trials of each class were shuffled before splitting in a pseudo-randomized manner. In each fold, the predictions of the classifier for the testing set were used to compute the Balanced Accuracy score and the F1-score of the classifier (see definition and method for calculation of these scores below). We then computed the mean Balanced Accuracy and F1 scores across folds, as well as their confidence interval. F1 scores can be found in Table S5.
- For the participants without narcolepsy, since *responsive* trials were scarce in particular during N2 sleep and REM sleep, we decided to train our classifier with the data of the participants with narcolepsy and to test its performance on data from participants without narcolepsy. Specifically, we fitted our classifier with the N2 sleep trials from participants with narcolepsy, and then tested its predictions on N2 sleep trials from the participants without narcolepsy. As before, we computed balanced accuracy and f1 scores. To obtain a distribution of scores in the absence of cross-validation, we repeated the fitting and testing steps 10 times (note that the random parameters of the Random Forest classifier allowed us to obtain a distribution of -closely related-scores in this manner).

As mentioned above, we computed two scores to measure the performance of our classifier, both measures being well adapted to unbalanced datasets^52^ as ours (with more non-responsive trials than responsive ones during sleep):

- The balanced accuracy score corresponds, in binary classification problems, to the mean of the *sensitivity* (Se) (“How many relevant items are retrieved?”) and the *specificity* (Sp) (“How many non-relevant items are correctly identified”). In terms of true positives (TP), false negatives (FN), true negatives (TN) and false positives (FP) (where, in our case, true positives are responsive trials correctly identified by the classifier, and true negatives non-responsive trials correctly identified by the classifier), the balanced accuracy score can be computed by the following formula:

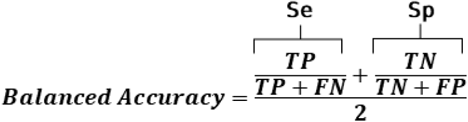
- The F1-score corresponds, in binary classification problems, to the harmonic mean of the *precision* (“How many retrieved items are relevant?”) and the *sensitivity*. It can be computed by the following formula:

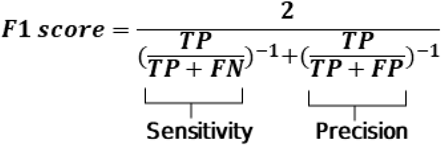

### Decoding of stimulus-related brain activity

We aimed at assessing brain responses to stimuli in function of participants’ sleep/wake stages and of their responsiveness to the task. Our unbalanced datasets across participants and conditions did not allow for a more classical approach such as ERP (Evoked Related Potentials). We hence decided to use a multivariate pattern analysis (MPVA) with the temporal generalization decoding method^31^. The idea of this analysis is to test, for a given time-point after stimulus presentation, how different the multivariate pattern of activity across electrodes was at this specific time point compared to the pattern at baseline (before stimulus presentation), for the different conditions.

To reduce computation time, we first down-sampled our data to 100Hz (decimation factor of 2.5). To ensure a correct features/number of trials ratio, we restricted our analysis to 3 centro-parietal electrodes (Cz, Pz, and P3), and, for each condition (sleep stage/responsiveness), we only included in our analysis the participants who had at least 15 trials of the given condition. Given these restrictions, we only had enough participants for statistical analysis for Lucid REM sleep (10 participants for responsive trials, and 9 participants for non-responsive trials) and for Wake (14 participants for responsive trials). Then, for each condition, participant, trial, and channel, we computed the mean voltage during the 350ms baseline period before stimulus presentation and used this value to create dummy “baseline” trials with the same dimensionality as the original trials. Note that after this step, for each condition and each participant, we obtained a balanced set of dummy “baseline trials” (reflecting baseline brain activity before/without stimulus presentation) and actual trials where the stimulus was presented.

Then, independently for each condition and each participant, we trained a linear classifier to decode stimulus-present versus stimulus-absent trials (“baseline” dummy trials versus actual trials), using an L2-regularized (C=1) logistic regression, in a 5-fold cross-validation procedure. In each fold, all the trials were shuffled in a pseudo-randomized manner and split into a training set (⅘of the trials) and a testing set (⅕of the trials). The features (channel amplitudes) were standardized across training trials before being provided to the classifier for training. This training procedure was applied at each time step independently. Following the time generalization approach, the model trained at each time step was then tested at all the time steps on the testing set trials, at each cross-validation fold. The classifier performance at each training and testing time was evaluated by the area under the receiver operating curve (AUC) at each cross-validation fold. At the end of the cross-validation procedure, the global performance of the classifier at each training and testing time was obtained by averaging the intermediate values obtained at each fold, for each participant and each experimental condition. Group-level performance for each condition was finally obtained by averaging across participants, independently for each condition (stage /responsiveness).

### Statistical analysis

Most statistical analyses were conducted in *R*^53^ using the *lme4*^54^, *emmeans*^55^, *BayesFactor*^56^ and *DHARMa*^57^ packages. For the machine learning analysis, statistics were conducted in *Python*^49^ using the *numpy*^58^, *scipy*^59^, and *scikit-learn*^60^ packages. All statistics were corrected for multiple comparisons using the False Discovery Rate (FDR) Benjamini–Hochberg procedure.

#### Behavior

Linear mixed models with subject ID as a random factor were used for all statistical analyses. We evaluated participants’ ability to respond to stimuli in different sleep stages (Figure 1A and Figure 2B). First, we focused on the comparison between the ON and OFF periods separately for each sleep stage. Binomial generalized linear mixed models with stimulation period (ON vs OFF) as the independent variable and responsiveness (response vs no response; both contraction types combined) as the dependent variable were used in this analysis. The models’ assumptions were evaluated using the DHARMa^57^ package in *R*^53^. Next, we focused on the “ON” stimulation periods during which participants were presented with stimuli. The model had sleep stages (wake, N1, N2, N3, REM sleep in healthy participants and wake, N1 N2, N3, non-lucid REM sleep and lucid REM sleep in participants with narcolepsy) as the independent variable and responsiveness (response vs no response) as the dependent variable. For accuracy, we computed the percentage of correct responses for each participant at each sleep stage and compared them to the 50% chance level using Wilcoxon signed rank test. Only participants with at least 3 responses are included in this analysis. Finally, the differences in reaction times in different sleep stages were assessed using a linear mixed model (Figure 1C). An inverse transformation was applied to the reaction times (1/RT) to better fit the model assumptions.

#### EEG markers

In order to investigate how different neural markers differ in trials with a response and without any response, we first z-scored marker values at subject level. We then used a mixed linear model for each EEG marker with subject ID as a random factor, responsiveness as the independent variable, and the EEG marker as the dependent variable. The analysis was conducted at a single-trial level. Since the responsiveness and the sleep stage are not independent (for example, in wake we observed more responses than in N2 sleep), we could not include the sleep stage as an additional independent variable in the models. Thus, we performed the tests separately for each sleep stage, resulting in a test for each marker in each sleep stage. We computed a similar analysis to compare, in REM sleep, lucid and non-lucid trials.

#### Prediction of responsiveness at a trial level using a Random Forest Classifier

We scored classifier performance at each sleep/wake stage and for each group using the Balanced Accuracy score and the F1-score (cf above). To assess how different these scores were from chance level, we performed, independently for each score, a 500-permutation procedure. At each permutation, trial labels (*responsive* versus *non-responsive*) were randomly shuffled, and the entire 10-fold cross-validation procedure was performed, allowing us to obtain a distribution of chance-level scores. To calculate the p-value for each state, we counted the number of permutation scores equal or higher to our true score and divided it by the number of permutations plus one.

#### Decoding of stimulus-related brain activity using temporal generalization decoding

For each experimental condition (sleep stage/responsiveness), classification performance at each training and testing time was tested against 0.5 (chance) using a two-sided non-parametric sign test across subjects, and these statistics were then corrected for multiple comparisons using the False Discovery Rate (FDR) Benjamini–Hochberg procedure. In figure 5 panel B, significant time points (p<0.05 FDR corrected) with an AUC>0.5 are outlined in black.

## Acknowledgments

This study was funded by the Agence Nationale de la Recherche (ANR-20-CE37-0001-01, grant to DO) and a research grant from Société Française de Recherche et Médecine du Sommeil (DO). BT and EMM received PhD grants from the French Ministry of Higher Education and the INSERM (Institut national de la santé et de la recherche médicale) while JBM received a grant from APHP and Sorbonne University (‘Poste d’accueil APHP’). The National Reference Center for Narcolepsy benefits from a recurrent grant from the National Program on Rare Diseases (PNMR-3 grant to IA) from the Health Ministry, which encourages research on narcolepsy.

Authors would like to thank Sana Rebbah for her help in the statistical analyses.

## Author contributions

Conceptualization: BT, EC, DO

Data collection: BT, EC, AFG, JBM

Data analysis: BT, EMM, EC, JBM, NW, PP

Visualization: BT, EMM, EC

Supervision: JS, IA, LN, DO

Writing: BT, EMM, EC, IA, LN, DO

## Data and materials availability

All the relevant data is available upon reasonable request. Inquiries should be directed to the corresponding author.

## Supplementary Materials

- Supplementary results
- Figures S1-S5
- Tables S1-S8

## References

1. Andrillon, T. & Kouider, S. The vigilant sleeper: neural mechanisms of sensory (de)coupling during sleep. Curr. Opin. Physiol. 15, 47–59 (2020).

2. Bastuji, H. & García-Larrea, L. Evoked potentials as a tool for the investigation of human sleep. Sleep Med. Rev. 3, 23–45 (1999).

3. Ruby, P., Caclin, A., Boulet, S., Delpuech, C. & Morlet, D. Odd sound processing in the sleeping brain. J. Cogn. Neurosci. 20, 296–311 (2008).

4. Strauss, M. et al. Disruption of hierarchical predictive coding during sleep. Proc. Natl. Acad. Sci. U. S. A. 112, E1353–1362 (2015).

5. Issa, E. B. & Wang, X. Altered Neural Responses to Sounds in Primate Primary Auditory Cortex during Slow-Wave Sleep. J. Neurosci. 31, 2965–2973 (2011).

6. Nir, Y., Vyazovskiy, V. V., Cirelli, C., Banks, M. I. & Tononi, G. Auditory Responses and Stimulus-Specific Adaptation in Rat Auditory Cortex are Preserved Across NREM and REM Sleep. Cereb. Cortex 25, 1362–1378 (2015).

7. Kouider, S., Andrillon, T., Barbosa, L. S., Goupil, L. & Bekinschtein, T. A. Inducing Task-Relevant Responses to Speech in the Sleeping Brain. Curr. Biol. 24, 2208–2214 (2014).

8. Oudiette, D. & Paller, K. A. Upgrading the sleeping brain with targeted memory reactivation. Trends Cogn. Sci. 17, 142–149 (2013).

9. Rudoy, J. D., Voss, J. L., Westerberg, C. E. & Paller, K. A. Strengthening individual memories by reactivating them during sleep. Science 326, 1079 (2009).

10. Rasch, B., Büchel, C., Gais, S. & Born, J. Odor Cues During Slow-Wave Sleep Prompt Declarative Memory Consolidation. Science 315, 1426–1429 (2007).

11. Arzi, A. et al. Olfactory Aversive Conditioning during Sleep Reduces Cigarette-Smoking Behavior. J. Neurosci. 34, 15382–15393 (2014).

12. Andrillon, T., Poulsen, A. T., Hansen, L. K., Léger, D. & Kouider, S. Neural Markers of Responsiveness to the Environment in Human Sleep. J. Neurosci. 36, 6583–6596 (2016).

13. Solomonova, E. & Carr, M. Incorporation of External Stimuli into Dream Content. in 213–218 (2019).

14. Konkoly, K. R. et al. Real-time dialogue between experimenters and dreamers during REM sleep. Curr. Biol. 31, 1417-1427.e6 (2021).

15. Strauss, M., Sitt, J. D., Naccache, L. & Raimondo, F. Predicting the loss of responsiveness when falling asleep in humans. NeuroImage 251, 119003 (2022).

16. Ogilvie, R. D. The process of falling asleep. Sleep Med. Rev. 5, 247–270 (2001).

17. Rivera-García, A. P., Ramírez-Salado, I., Corsi-Cabrera, M. & Calvo, J. M. Facial muscle activation during sleep and its relation to the rapid eye movements of REM sleep. J. Sleep Res. 20, 82–91 (2011).

18. LaBerge, S. P., Nagel, L. E., Dement, W. C. & Zarcone, V. P. Lucid dreaming verified by volitional communication during REM sleep. Percept. Mot. Skills 52, 727–732 (1981).

19. Dodet, P., Chavez, M., Leu-Semenescu, S., Golmard, J.-L. & Arnulf, I. Lucid Dreaming in Narcolepsy. Sleep 38, 487–497 (2015).

20. Berry, R. B. et al. AASM Scoring Manual Updates for 2017 (Version 2.4). J. Clin. Sleep Med. JCSM Off. Publ. Am. Acad. Sleep Med. 13, 665–666 (2017).

21. Perea, M., Marcet, A., Vergara-Martínez, M. & Gomez, P. On the Dissociation of Word/Nonword Repetition Effects in Lexical Decision: An Evidence Accumulation Account. Front. Psychol. 7, (2016).

22. Sitt, J. D. et al. Large scale screening of neural signatures of consciousness in patients in a vegetative or minimally conscious state. Brain 137, 2258–2270 (2014).

23. Martínez Vivot, R., Pallavicini, C., Zamberlan, F., Vigo, D. & Tagliazucchi, E. Meditation Increases the Entropy of Brain Oscillatory Activity. Neuroscience 431, 40–51 (2020).

24. King, J.-R. et al. Information Sharing in the Brain Indexes Consciousness in Noncommunicative Patients. Curr. Biol. 23, 1914–1919 (2013).

25. Engemann, D. A. et al. Robust EEG-based cross-site and cross-protocol classification of states of consciousness. Brain 141, 3179–3192 (2018).

26. Bourdillon, P. et al. Brain-scale cortico-cortical functional connectivity in the delta-theta band is a robust signature of conscious states: an intracranial and scalp EEG study. Sci. Rep. 10, 14037 (2020).

27. Lee, M. D. & Wagenmakers, E.-J. Bayesian Cognitive Modeling: A Practical Course. (Cambridge University Press, 2014).

28. Del Cul, A., Baillet, S. & Dehaene, S. Brain dynamics underlying the nonlinear threshold for access to consciousness. PLoS Biol. 5, e260 (2007).

29. Bekinschtein, T. A. et al. Neural signature of the conscious processing of auditory regularities. Proc. Natl. Acad. Sci. 106, 1672–1677 (2009).

30. Sergent, C., Baillet, S. & Dehaene, S. Timing of the brain events underlying access to consciousness during the attentional blink. Nat. Neurosci. 8, 1391–1400 (2005).

31. King, J.-R. & Dehaene, S. Characterizing the dynamics of mental representations: the temporal generalization method. Trends Cogn. Sci. 18, 203–210 (2014).

32. Sanchez, G., Hartmann, T., Fuscà, M., Demarchi, G. & Weisz, N. Decoding across sensory modalities reveals common supramodal signatures of conscious perception. Proc. Natl. Acad. Sci. 117, 7437–7446 (2020).

33. Sergent, C. et al. Bifurcation in brain dynamics reveals a signature of conscious processing independent of report. Nat. Commun. 12, 1149 (2021).

34. Oudiette, D. et al. REM sleep respiratory behaviours match mental content in narcoleptic lucid dreamers. Sci. Rep. 8, 2636 (2018).

35. Voss, U., Holzmann, R., Tuin, I. & Hobson, J. A. Lucid dreaming: a state of consciousness with features of both waking and non-lucid dreaming. Sleep 32, 1191–1200 (2009).

36. Andrillon, T. et al. Does the Mind Wander When the Brain Takes a Break? Local Sleep in Wakefulness, Attentional Lapses and Mind-Wandering. Front. Neurosci. 13, (2019).

37. Andrillon, T., Burns, A., Mackay, T., Windt, J. & Tsuchiya, N. Predicting lapses of attention with sleep-like slow waves. Nat. Commun. 12, 3657 (2021).

38. Dauvilliers, Y. et al. REM Sleep Characteristics in Narcolepsy and REM Sleep Behavior Disorder. Sleep 30, 844–849 (2007).

39. Dresler, M. et al. Neural correlates of dream lucidity obtained from contrasting lucid versus non-lucid REM sleep: a combined EEG/fMRI case study. Sleep 35, 1017–1020 (2012).

40. Baird, B., Tononi, G. & LaBerge, S. Lucid dreaming occurs in activated rapid eye movement sleep, not a mixture of sleep and wakefulness. Sleep zsab294 (2022) doi:10.1093/sleep/zsab294.

41. Filevich, E., Dresler, M., Brick, T. R. & Kühn, S. Metacognitive Mechanisms Underlying Lucid Dreaming. J. Neurosci. 35, 1082–1088 (2015).

42. Dresler, M. et al. Volitional components of consciousness vary across wakefulness, dreaming and lucid dreaming. Front. Psychol. 4, (2014).

43. Rossetti, Y. Implicit short-lived motor representations of space in brain damaged and healthy subjects. Conscious. Cogn. 7, 520–558 (1998).

44. Stumbrys, T. & Erlacher, D. Lucid dreaming during NREM sleep: Two case reports. Int. J. Dream Res. 5, 151–155 (2012).

45. Ferrand, L. et al. MEGALEX: A megastudy of visual and auditory word recognition. Behav. Res. Methods 50, 1285–1307 (2018).

46. Naccache, L. Why and how access consciousness can account for phenomenal consciousness. Philos. Trans. R. Soc. Lond. B. Biol. Sci. 373, 20170357 (2018).

47. Brainard, D. H. The Psychophysics Toolbox. Spat. Vis. 10, 433–436 (1997).

48. Jas, M., Engemann, D. A., Bekhti, Y., Raimondo, F. & Gramfort, A. Autoreject: Automated artifact rejection for MEG and EEG data. NeuroImage 159, 417–429 (2017).

49. Van Rossum, G. & Drake, F. L. Python 3 Reference Manual. (CreateSpace, 2009).

50. Richman, J. S. & Moorman, J. R. Physiological time-series analysis using approximate entropy and sample entropy. Am. J. Physiol. Heart Circ. Physiol. 278, H2039–2049 (2000).

51. Fernandez-Delgado, M., Cernadas, E., Barro, S. & Amorim, D. Do we Need Hundreds of Classifiers to Solve Real World Classification Problems? J. Mach. Learn. Res. 15, 3133–3181 (2014).

52. Kelleher, J. D., Namee, B. M. & D’Arcy, A. Fundamentals of Machine Learning for Predictive Data Analytics: Algorithms, Worked Examples, and Case Studies. (MIT Press, 2015).

53. R Core Team. R: A Language and Environment for Statistical Computing. (R Foundation for Statistical Computing, 2021).

54. Bates, D., Mächler, M., Bolker, B. & Walker, S. Fitting Linear Mixed-Effects Models Using lme4. J. Stat. Softw. 67, 1–48 (2015).

55. Lenth, R. V. emmeans: Estimated Marginal Means, aka Least-Squares Means. R package version 1.6.2-1. (2021).

56. Morey, R. D. & Rouder, J. N. BayesFactor. (2013).

57. Hartig, F. DHARMa: Residual Diagnostics for Hierarchical (Multi-Level / Mixed) Regression Models. (2022).

58. Harris, C. R. et al. Array programming with NumPy. Nature 585, 357–362 (2020).

59. Virtanen, P. et al. SciPy 1.0: fundamental algorithms for scientific computing in Python. Nat. Methods 17, 261–272 (2020).

60. Pedregosa, F. et al. Scikit-learn: Machine Learning in Python. J. Mach. Learn. Res. 12, 2825– 2830 (2011).

