## Supplemental Data for "Behavioral and brain responses to verbal stimuli reveal transient periods of cognitive integration of external world in all sleep stages"

[4] AP-HP, Hôpital Pitié-Salpêtrière, Service de Pneumologie, Médecine Intensive et Réanimation (Département R3S), Paris 75013, France

### SUPPLEMENTARY MATERIALS

#### SUPPLEMENTARY RESULTS

##### Mixed contractions to signal lucidity

In this experiment, participants were instructed to perform a lexical decision task in their sleep by contracting 3 times their corrugator or zygomatic muscles (according to the stimulus). In case they were lucid dreaming but were not hearing any stimuli, they were asked to signal their lucidity with a “mixed contraction” by alternating one corrugator and one zygomatic muscle contraction. We did not observe any mixed contractions in participants without narcolepsy (HP). On the other hand, we observed a total of 117 mixed contractions from 12 participants with narcolepsy (NP) in 19 different naps. Importantly, all 19 naps contained responses to the stimuli during N2 and/or REM sleep. Among the 117 mixed contractions, 93 were observed in REM sleep, 92 being in naps that were reported (upon awakening) to be lucid. Moreover, 18 contractions were observed in N2 sleep (12 being in lucid naps) and 6 contractions were observed in N1 sleep (5 being in lucid naps). These results indicate that (I) facial-muscle contractions can be used to signal lucidity, validating our previously published results[14]; (II) participants remember most of the signaled lucidity episodes, especially in REM sleep; and (III) all signaled lucidity episodes are associated with higher responsiveness to the external stimuli.

##### Old/New recognition task upon awakening

After each nap, participants performed an old-new recognition task, during which they were presented with stimuli they heard during the preceding nap and new stimuli that were never presented during the experiment. Participants had to indicate whether they had heard the stimuli during the preceding session with one of the following responses: 1: I heard it from the dream (for example, a person from their dream saying the word), 2: I heard it from outside (pronounced by the computer), 3: I am not sure I heard it, 4: I am sure I did not hear it. They responded by pressing the corresponding button without any time pressure. The four options were explained to the participants during training, prior to the first session. We assessed whether participants were able to correctly recognize the stimuli upon awakening. We focused on participants with narcolepsy (NP) since they went through 5 short naps (rather than a long 100 mins nap in HP) that would favor the recall. First, we computed, for each nap, the percentage of false recall of new stimuli. A stimulus was considered “recalled” if participants reported either (1) hearing it in their dreams or (2) hearing it from outside of their dreams while asleep. On average 8.46% of new stimuli were falsely recalled. Then, we assessed the correct recall of stimuli that were previously presented in different sleep stages. The percentage of correct recall was 21% in Wake, 16.26% in N1, 8.7% in N2, 9.1% in REM, 8.9% in lucid N2 and 11.1% in lucid REM sleep, significantly different than false recall only in Wake ($p < 0.0001$, $z = 4.16$) and N1 sleep ($p < 0.002$, $z = 3.7$).

##### Random Forest classification

All balanced accuracy scores were significantly different than the chance level computed by a 500-permutation procedure ($p = .002$ for all stages in NP, and $p = .004$ for N2 sleep in HP), with a mean balanced accuracy score of permutation trials very close to 50% for all stages. Similarly, f1 scores were significantly different than chance level ($p = 0.002$) for Wake, N2, and REM sleep in NP (with a statistical trend for N1, $p = 0.06$), and for N2 in HP (see Figure S4).

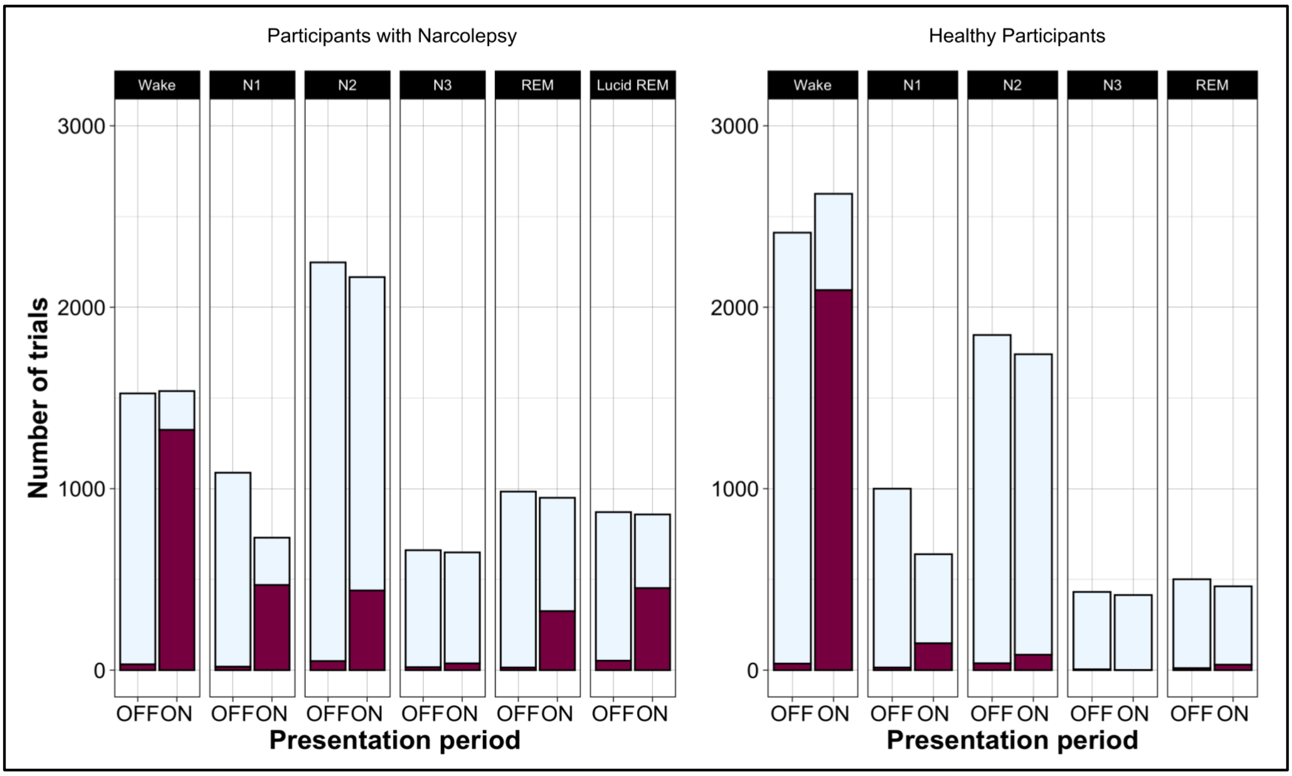


**Figure S1. Number of trials during ON and OFF stimulation periods in different sleep stages in participants with (left) and without (right) narcolepsy.** The partition of trials containing a response is filled with dark red color in both stimulation periods.

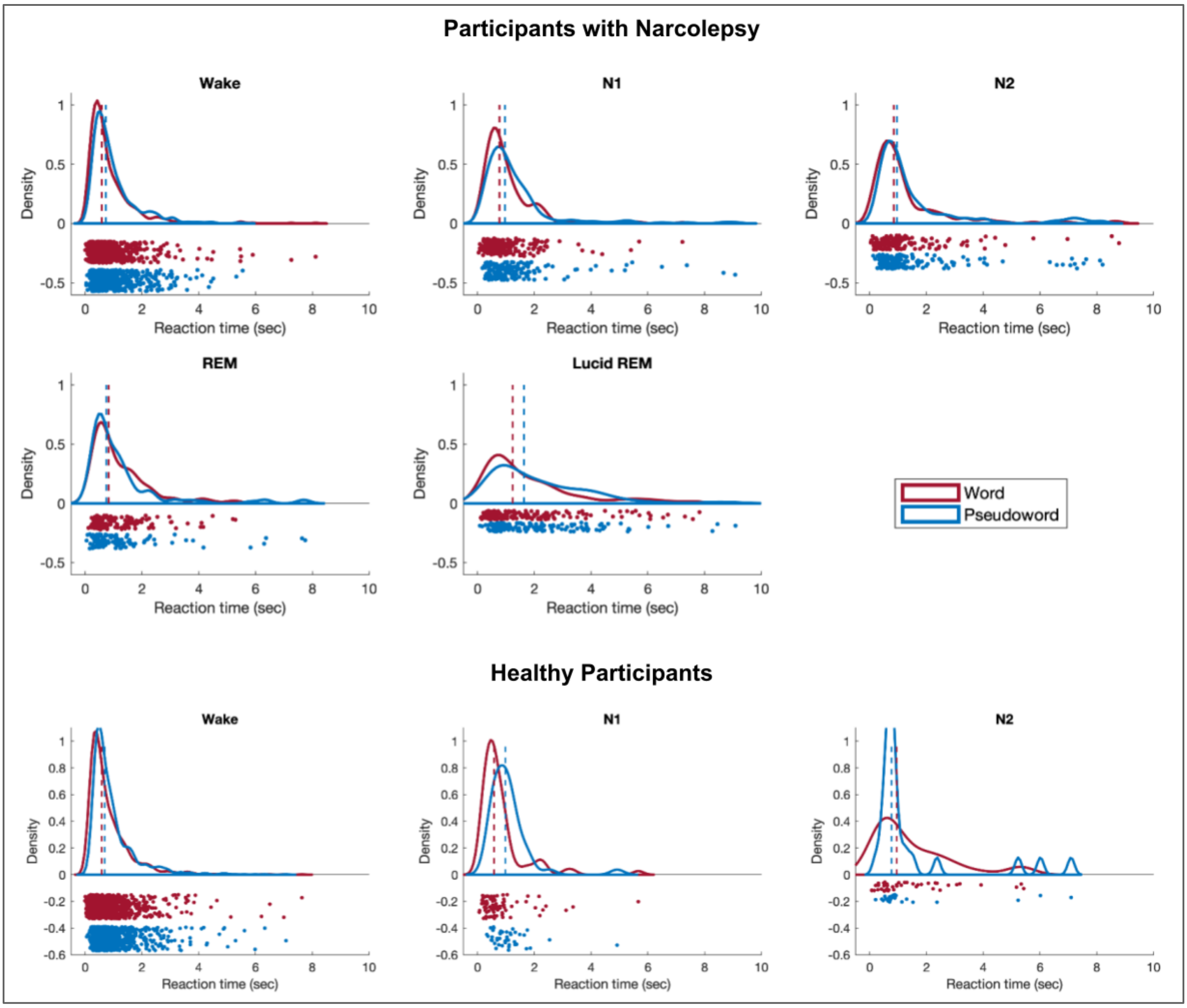


**Figure S2. Reaction times for words and pseudo-words in the different sleep/wake stages in participants with narcolepsy and healthy participants**

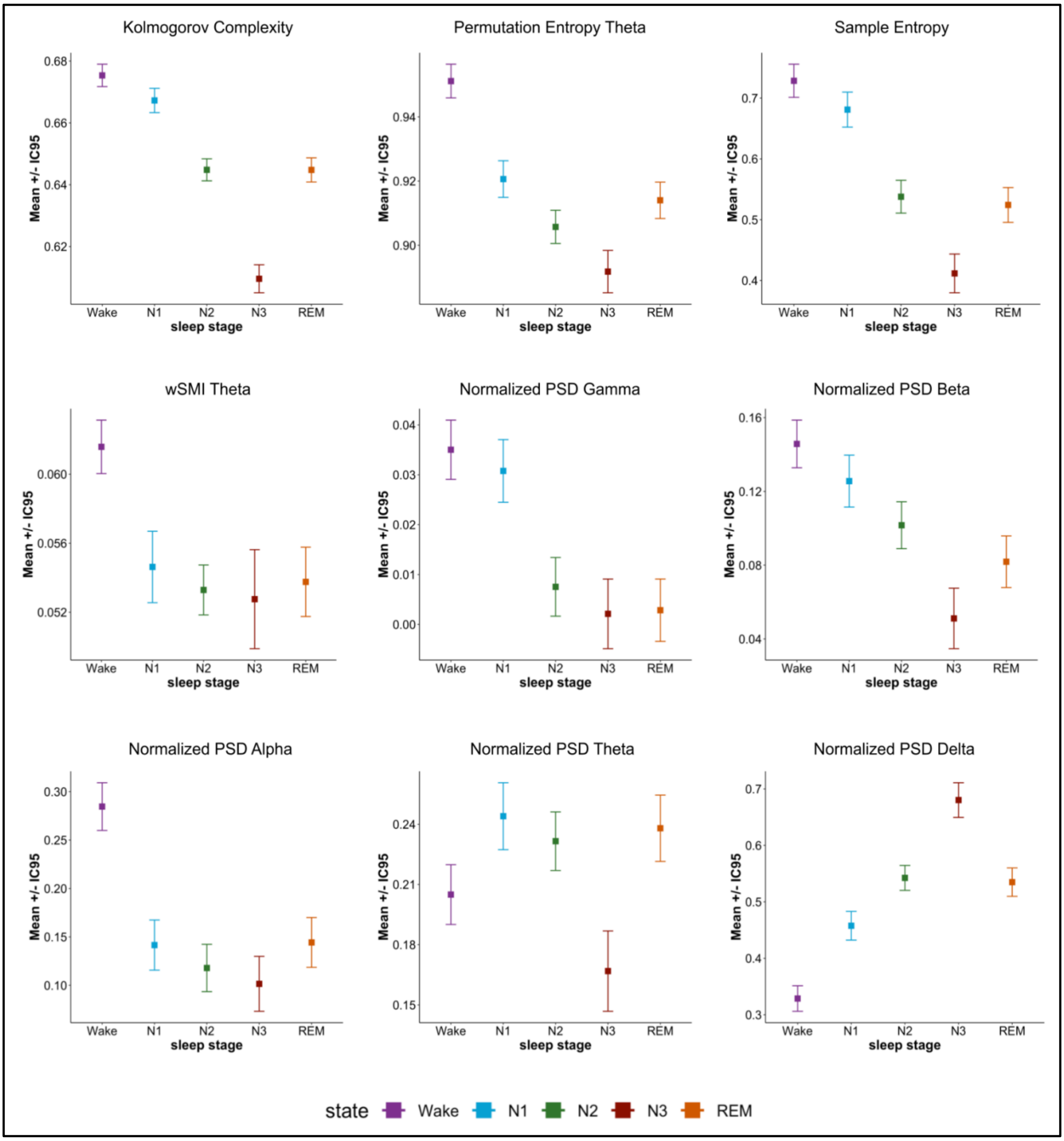


**Figure S3. Evolution of electrophysiological markers across sleep stages in HP.** Three complexity measures (the Kolmogorov Complexity -KC, the Permutation Entropy -PE, and the Sample Entropy -SE), one connectivity measure (weighted symbolic mutual information (wSMI) in the theta band), and five spectral measures (normalized power spectral densities (PSD) of delta, theta, alpha, beta and gamma frequency bands) were computed separately for the wake, N1, N2, N3, and REM sleep stages in HP. Each dot indicates marginal means estimated by a mixed-linear model including sleep stage as an independent variable, marker as the dependent variable, and participant ID as a random variable. Error bars denote 95% confidence intervals. Complexity and high-frequency PSD decreased in sleep compared to wake (wake > N1 > N2 ≈ REM > N3), whereas delta PSD increased with sleep (N3 > N2 ≈ REM > N1 > wake). Theta PSD was higher in N1 and lower in N3 sleep. Details of the statistical comparisons can be found in Table S3.

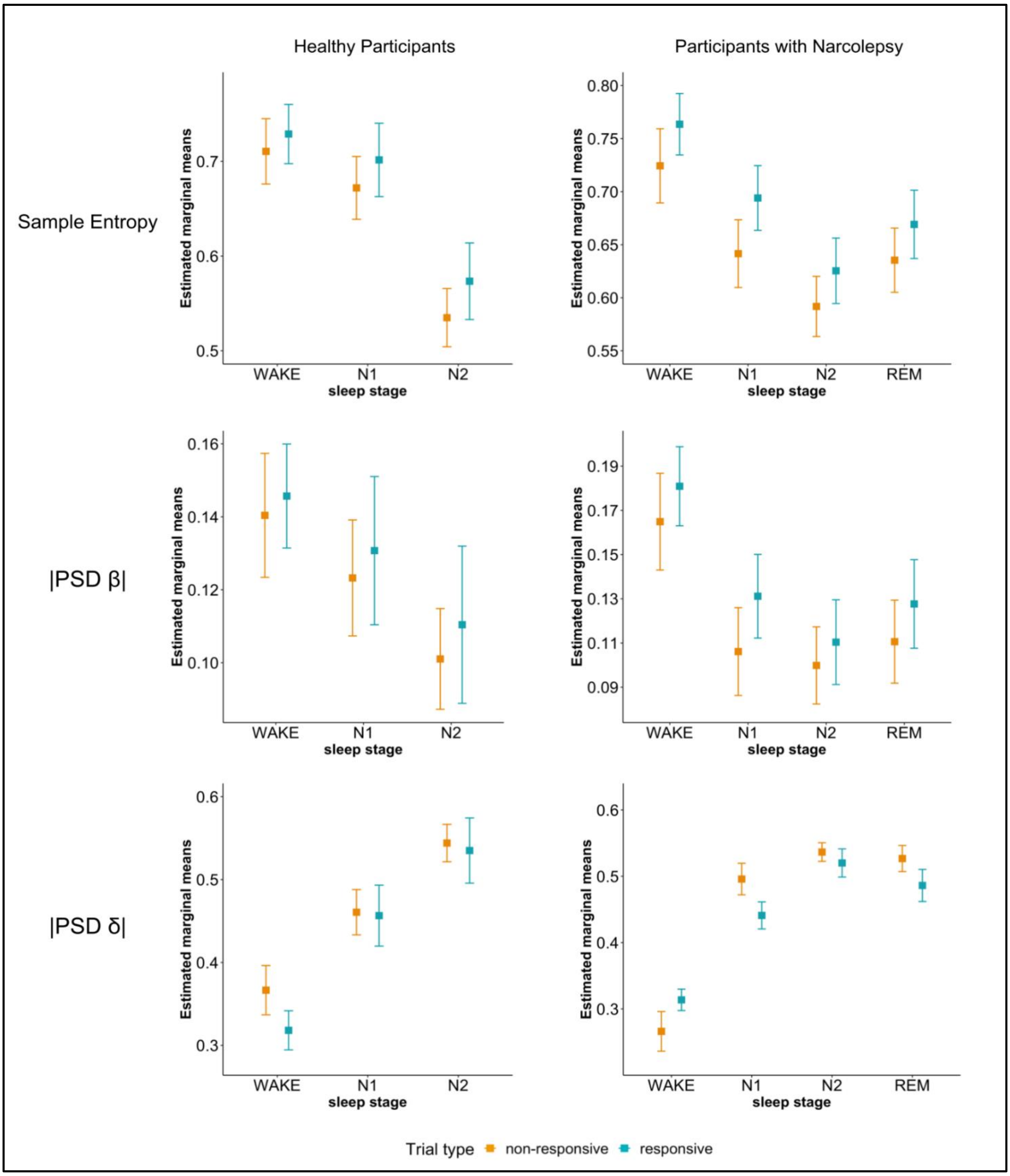


**Figure S4. Variations in sample entropy, normalized PSD in beta (|PSD β|) and delta (|PSD δ|) frequencies with responsiveness across different sleep stages in participants without (left) and with (right) narcolepsy.** Please note that marker values in different sleep stages were never at wake level, indicating that participants were indeed asleep while they were responding.

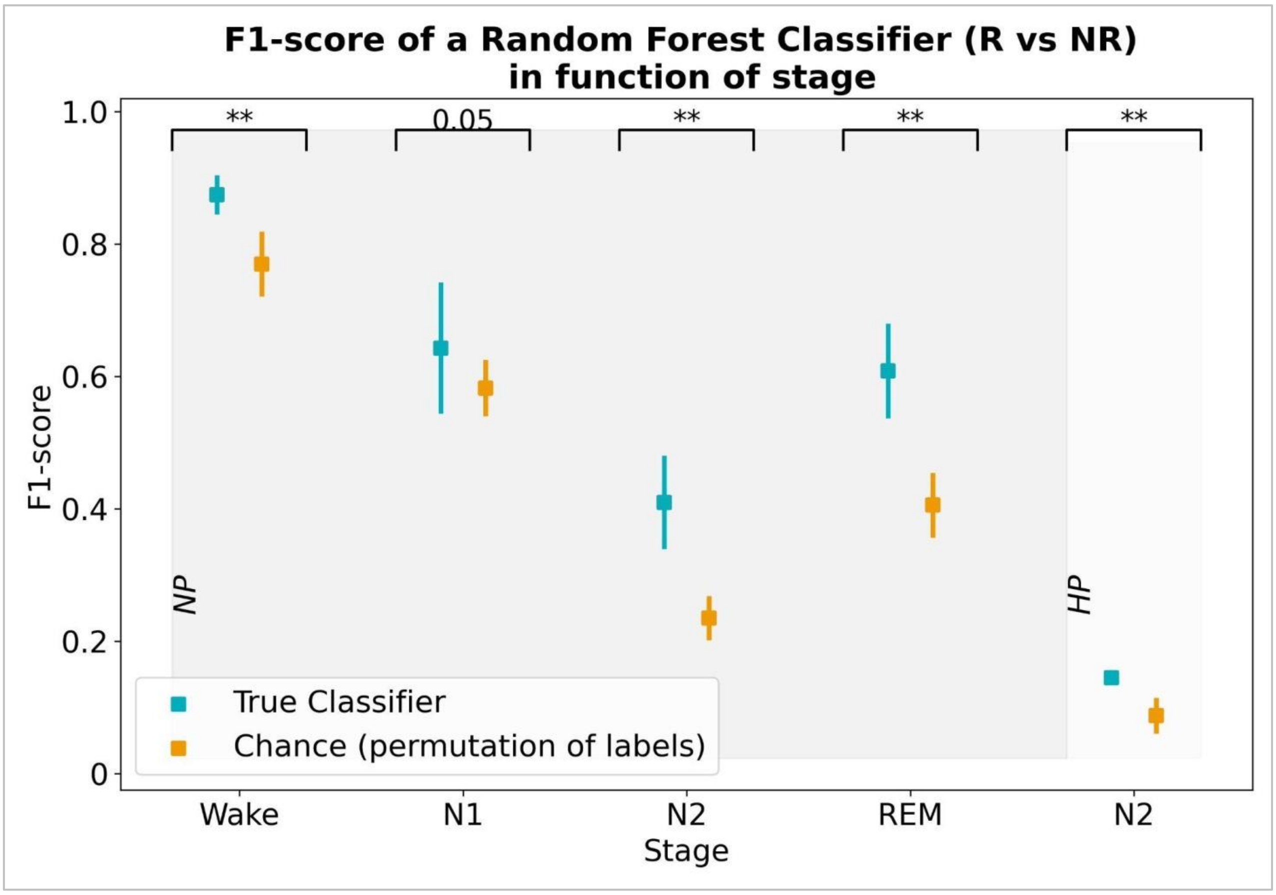


**Figure S5. F1-score of the Random Forest classifiers trained with the neurophysiological markers to classify between responsive vs non-responsive trials, in different sleep stages, for participants with narcolepsy (NP) and healthy participants (HP).** Blue (true classifier): true performance of the classifier (mean F1 score and 95IC across folds). Orange (Chance): chance-level performance computed with 500 random permutations of the data labels (mean F1 score and 95IC across permutations). Statistical difference between true performance and chance is showed for each sleep stage. **: $p<0.01$.

**Table S1. Details of the multiple comparisons of the response rates during ON periods in different sleep stages.**

| COMPARISON | Healthy Participants (HP) | | Participants with Narcolepsy (NP) | |
|---|---|---|---|---|
| | *z* | FDR corrected p-value | *z* | FDR corrected p-value |
| **Wake - N1** | 21.50 | < 0.0001 | 13.47 | < 0.0001 |
| **Wake - N2** | 33.23 | < 0.0001 | 31.26 | < 0.0001 |
| **Wake - N3** | 8.23 | < 0.0001 | 18.20 | < 0.0001 |
| **Wake - REM** | 19.32 | < 0.0001 | 21.89 | < 0.0001 |
| **Wake - Lucid REM** | - | - | 15.27 | < 0.0001 |
| **N1 - N2** | 12.56 | < 0.0001 | 19.51 | < 0.0001 |
| **N1 - N3** | 5.39 | < 0.0001 | 12.08 | < 0.0001 |
| **N1 - REM** | 6.42 | < 0.0001 | 9.57 | < 0.0001 |
| **N1 - Lucid REM** | - | - | 2.15 | 0.03 |
| **N2 - N3** | 3.39 | 0.0008 | 4.36 | < 0.0001 |
| **N2 - REM** | -2.42 | 0.01 | -8.54 | < 0.0001 |
| **N2 - Lucid REM** | - | - | -18.11 | < 0.0001 |
| **N3 - REM** | -3.90 | 0.0001 | -7.55 | < 0.0001 |
| **N3 - Lucid REM** | - | - | -11.11 | < 0.0001 |
| **REM - Lucid REM** | - | - | -7.87 | < 0.0001 |

**Table S2. Details of the multiple comparisons of neurophysiological markers in different sleep stages in NP.** K for Kolmogorov Complexity; PE θ for Permutation Entropy in the theta band; SE for Sample Entropy; and wSMI θ for weighted symbolic mutual information in the theta band. All p-values are FDR corrected for multiple comparisons. Statistically significative comparisons ($p < 0.05$) are highlighted.

| | K | PE θ | SE | wSMI θ | PSD \|γ\| | PSD \|β\| | PSD \|α\| | PSD \|θ\| | PSD \|δ\| |
|---|---|---|---|---|---|---|---|---|---|
| **Wake - N1** | $t$ = 9.88<br>$p$ < .0001 | $t$ = 21.08<br>$p$ < .0001 | $t$ = 12.37<br>$p$ < .0001 | $t$ = 6.29<br>$p$ < .0001 | $t$ = 9.54<br>$p$ < .0001 | $t$ = 13.63<br>$p$ < .0001 | $t$ = 25.91<br>$p$ < .0001 | $t$ = -12.44<br>$p$ < .0001 | $t$ = -17.95<br>$p$ < .0001 |
| **Wake -N2** | $t$ = 27.01<br>$p$ < .0001 | $t$ = 33.30<br>$p$ < .0001 | $t$ = 28.90<br>$p$ < .0001 | $t$ = 8.90<br>$p$ < .0001 | $t$ = 17.76<br>$p$ < .0001 | $t$ = 22.08<br>$p$ < .0001 | $t$ = 38.66<br>$p$ < .0001 | $t$ = -12.38<br>$p$ < .0001 | $t$ = -32.48<br>$p$ < .0001 |
| **Wake - N3** | $t$ = 44.86<br>$p$ < .0001 | $t$ = 33.84<br>$p$ < .0001 | $t$ = 39.14<br>$p$ < .0001 | $t$ = 6.24<br>$p$ < .0001 | $t$ = 19.70<br>$p$ < .0001 | $t$ = 26.95<br>$p$ < .0001 | $t$ = 31.16<br>$p$ < .0001 | $t$ = -1.53<br>$p$ = 0.14 | $t$ = -37.69<br>$p$ < .0001 |
| **Wake - REM** | $t$ = 13.63<br>$p$ < .0001 | $t$ = 24.95<br>$p$ < .0001 | $t$ = 15.78<br>$p$ < .0001 | $t$ = 8.08<br>$p$ < .0001 | $t$ = 10.35<br>$p$ < .0001 | $t$ = 14.35<br>$p$ < .0001 | $t$ = 31.50<br>$p$ < .0001 | $t$ = -10.51<br>$p$ < .0001 | $t$ = -23.44<br>$p$ < .0001 |
| **N1 - N2** | $t$ = 13.30<br>$p$ < .0001 | $t$ = 6.62<br>$p$ < .0001 | $t$ = 12.28<br>$p$ < .0001 | $t$ = 1.06<br>$p$ = 0.31 | $t$ = 5.41<br>$p$ < .0001 | $t$ = 4.82<br>$p$ < .0001 | $t$ = 6.18<br>$p$ < .0001 | $t$ = 2.55<br>$p$ = 0.13 | $t$ = -9.12<br>$p$ < .0001 |
| **N1 - N3** | $t$ = 34.63<br>$p$ < .0001 | $t$ = 14.63<br>$p$ < .0001 | $t$ = 27.09<br>$p$ < .0001 | $t$ = 0.68<br>$p$ = 0.51 | $t$ = 10.83<br>$p$ < .0001 | $t$ = 14.36<br>$p$ < .0001 | $t$ = 8.07<br>$p$ < .0001 | $t$ = 8.98<br>$p$ < .0001 | $t$ = -20.72<br>$p$ < .0001 |
| **N1 - REM** | $t$ = 3.55<br>$p$ = .0005 | $t$ = 3.83<br>$p$ = .0002 | $t$ = 3.29<br>$p$ = 0.001 | $t$ = 1.71<br>$p$ = 0.1 | $t$ = 0.90<br>$p$ = 0.39 | $t$ = 0.88<br>$p$ = 0.39 | $t$ = 5.48<br>$p$ < .0001 | $t$ = -1.53<br>$p$ = 0.14 | $t$ = -5.16<br>$p$ < .0001 |
| **N2 - N3** | $t$ = 29.05<br>$p$ < .0001 | $t$ = 11.36<br>$p$ < .0001 | $t$ = 21.00<br>$p$ < .0001 | $t$ = -0.15<br>$p$ = 0.88 | $t$ = 7.93<br>$p$ < .0001 | $t$ = 12.68<br>$p$ < .0001 | $t$ = 3.95<br>$p$ < .0001 | $t$ = 8.35<br>$p$< .0001 | $t$ = -16.32<br>$p$ < .0001 |
| **N2 - REM** | $t$ = -8.91<br>$p$ < .0001 | $t$ = -1.89<br>$p$ = 0.06 | $t$ = -8.20<br>$p$ < .0001 | $t$ = 1.06<br>$p$ = 0.31 | $t$ = -4.31<br>$p$ < .0001 | $t$ = -3.73<br>$p$ = 0.0002 | $t$ = 0.59<br>$p$ = 0.56 | $t$ = -0.66<br>$p$ = 0.52 | $t$ = 2.79<br>$p$ = 0.006 |
| **N3 - REM** | $t$ = -29.82<br>$p$ < .0001 | $t$ = -10.51<br>$p$ < .0001 | $t$ = -22.85<br>$p$ < .0001 | $t$ = 0.90<br>$p$ = 0.39 | $t$ = -9.53<br>$p$ < .0001 | $t$ = -12.90<br>$p$ < .0001 | $t$ = -2.73<br>$p$ = 0.007 | $t$ = -7.18<br>$p$ < .0001 | $t$ = 15.23<br>$p$ < .0001 |

**Table S3. Details of the multiple comparisons of neurophysiological markers in different sleep stages in HP.** K for Kolmogorov Complexity; PE θ for Permuation Entropy in the theta band; SE for Sample Entropy; and wSMI θ for weighted symbolic mutual information in the theta band. All p-values are FDR corrected for multiple comparisons. Statistically significative comparisons ($p$ < 0.05) are highlighted.

| | **K** | **PE θ** | **SE** | **wSMI θ** | **PSD \|γ\|** | **PSD \|β\|** | **PSD \|α\|** | **PSD \|θ\|** | **PSD \|δ\|** |
|---|---|---|---|---|---|---|---|---|---|
| **Wake - N1** | $t = 6.66$<br>$p < .0001$ | $t = 17.12$<br>$p < .0001$ | $t = 6.37$<br>$p < .0001$ | $t = 6.59$<br>$p < .0001$ | $t = 2.56$<br>$p = 0.013$ | $t = 4.47$<br>$p < .0001$ | $t = 22.52$<br>$p < .0001$ | $t = -6.58$<br>$p < .0001$ | $t = -13.93$<br>$p < .0001$ |
| **Wake -N2** | $t = 33.86$<br>$p < .0001$ | $t = 34.39$<br>$p < .0001$ | $t = 34.53$<br>$p < .0001$ | $t = 10.79$<br>$p < .0001$ | $t = 22.36$<br>$p < .0001$ | $t = 13.18$<br>$p < .0001$ | $t = 25.37$<br>$p < .0001$ | $t = -6.06$<br>$p < .0001$ | $t = -31.23$<br>$p < .0001$ |
| **Wake - N3** | $t = 38.01$<br>$p < .0001$ | $t = 23.42$<br>$p < .0001$ | $t = 29.93$<br>$p < .0001$ | $t = 5.92$<br>$p < .0001$ | $t = 13.96$<br>$p < .0001$ | $t = 14.74$<br>$p < .0001$ | $t = 20.26$<br>$p < .0001$ | $t = 4.52$<br>$p < .0001$ | $t = -26.74$<br>$p < .0001$ |
| **Wake - REM** | $t = 24.34$<br>$p < .0001$ | $t = 20.19$<br>$p < .0001$ | $t = 26.56$<br>$p < .0001$ | $t = 7.34$<br>$p < .0001$ | $t = 18.79$<br>$p < .0001$ | $t = 13.72$<br>$p < .0001$ | $t = 21.37$<br>$p < .0001$ | $t = -5.41$<br>$p < .0001$ | $t = -21.65$<br>$p < .0001$ |
| **N1 - N2** | $t = 18.67$<br>$p < .0001$ | $t = 8.46$<br>$p < .0001$ | $t = 19.50$<br>$p < .0001$ | $t = 1.29$<br>$p = 0.22$ | $t = 14.20$<br>$p < .0001$ | $t = 5.36$<br>$p < .0001$ | $t = 3.77$<br>$p = 0.0002$ | $t = 2.13$<br>$p = 0.04$ | $t = -9.28$<br>$p < .0001$ |
| **N1 - N3** | $t = 30.44$<br>$p < .0001$ | $t = 10.39$<br>$p < .0001$ | $t = 23.25$<br>$p < .0001$ | $t = 1.14$<br>$p = 0.28$ | $t = 11.10$<br>$p < .0001$ | $t = 10.59$<br>$p < .0001$ | $t = 4.05$<br>$p < .0001$ | $t = 8.35$<br>$p < .0001$ | $t = -15.46$<br>$p < .0001$ |
| **N1 - REM** | $t = 15.32$<br>$p = .0009$ | $t = 3.08$<br>$p = 0.003$ | $t = 17.48$<br>$p < .0001$ | $t = 0.69$<br>$p = 0.53$ | $t = 13.98$<br>$p < .0001$ | $t = 8.03$<br>$p < .0001$ | $t = -0.36$<br>$p = 0.73$ | $t = 0.84$<br>$p = 0.43$ | $t = -6.93$<br>$p < .0001$ |
| **N2 - N3** | $t = 21.20$<br>$p < .0001$ | $t = 5.72$<br>$p < .0001$ | $t = 12.41$<br>$p < .0001$ | $t = 0.37$<br>$p = 0.74$ | $t = 2.39$<br>$p = 0.02$ | $t = 8.20$<br>$p < .0001$ | $t = 1.89$<br>$p = 0.07$ | $t = 7.99$<br>$p < .0001$ | $t = -10.91$<br>$p < .0001$ |
| **N2 - REM** | $t = 0.03$<br>$p = 0.98$ | $t = -5.00$<br>$p < .0001$ | $t = 1.96$<br>$p = 0.06$ | $t = -0.48$<br>$p = 0.66$ | $t = 3.03$<br>$p = 0.003$ | $t = 4.72$<br>$p < .0001$ | $t = -4.46$<br>$p < .0001$ | $t = -1.17$<br>$p = 0.27$ | $t = 0.87$<br>$p = 0.42$ |
| **N3 - REM** | $t = -19.41$<br>$p < .0001$ | $t = -8.36$<br>$p < .0001$ | $t = -10.15$<br>$p < .0001$ | $t = -0.64$<br>$p = 0.55$ | $t = -0.29$<br>$p = 0.78$ | $t = -4.57$<br>$p < .0001$ | $t = -4.51$<br>$p < .0001$ | $t = -8.05$<br>$p < .0001$ | $t = 10.54$<br>$p < .0001$ |

**Table S4 Statistical differences of the neurophysiological markers between responsive and non-responsive trials in different sleep stages, in non-lucid naps, for HP and NP.** K for Kolmogorov Complexity; PE θ for Permutation Entropy in the theta band; SE for Sample Entropy; and wSMI θ for weighted symbolic mutual information in the theta band. All p-values are FDR corrected for multiple comparisons. Statistically significant comparisons ($p < 0.05$) are highlighted.

| **Participants with Narcolepsy (NP)** | | | | | | | | | |
|---|---|---|---|---|---|---|---|---|---|
| | **K** | **PE θ** | **SE** | **wSMI θ** | **PSD \|γ\|** | **PSD \|β\|** | **PSD \|α\|** | **PSD \|θ\|** | **PSD \|δ\|** |
| **Wake** | *t* = 2.63<br>*p* = 0.024 | *t* = 0.14<br>*p* = 0.96 | *t* = 3.15<br>*p* = 0.007 | *t* = -0.22<br>*p* = 0.93 | *t* = 2.55<br>*p* = 0.025 | *t* = 0.84<br>*p* = 0.53 | *t* = 0.65<br>*p* = 0.66 | *t* = -0.07<br>*p* = 0.99 | *t* = -1.18<br>*p* = 0.35 |
| **N1** | *t* = 4.52<br>*p* < .0001 | *t* = 1.48<br>*p* = 0.21 | *t* = 3.87<br>*p* = 0.0006 | *t* = -0.01<br>*p* = 0.99 | *t* = 2.60<br>*p* = 0.025 | *t* = 2.57<br>*p* = 0.025 | *t* = 2.30<br>*p* = 0.043 | *t* = -0.63<br>*p* = 0.66 | *t* = -4<br>*p* = 0.0005 |
| **N2** | *t* = 4.94<br>*p* < .0001 | *t* = 1.85<br>*p* = 0.11 | *t* = 5.03<br>*p* < .0001 | *t* = -0.51<br>*p* = 0.71 | *t* = 5.32<br>*p* < .0001 | *t* = 2.28<br>*p* = 0.043 | *t* = 0.56<br>*p* = 0.69 | *t* = -0.86<br>*p* = 0.53 | *t* = -1.47<br>*p* = 0.21 |
| **REM** | *t* = 3.91<br>*p* = 0.0006 | *t* = 2.63<br>*p* = 0.02 | *t* = 3.5<br>*p* = 0.002 | *t* = 1.62<br>*p* = 0.18 | *t* = 2.34<br>*p* = 0.041 | *t* = 2.94<br>*p* = 0.012 | *t* = 1.87<br>*p* = 0.11 | *t* = 0.05<br>*p* = 0.99 | *t* = -2.93<br>*p* = 0.012 |
| **Healthy Participants (HP)** | | | | | | | | | |
| | **K** | **PE θ** | **SE** | **wSMI θ** | **PSD \|γ\|** | **PSD \|β\|** | **PSD \|α\|** | **PSD \|θ\|** | **PSD \|δ\|** |
| **Wake** | *t* = 2.42<br>*p* = 0.048 | *t* = 3.78<br>*p* = 0.005 | *t* = 1.26<br>*p* = 0.40 | *t* = -0.46<br>*p* = 0.76 | *t* = 0.21<br>*p* = 0.87 | *t* = 2.32<br>*p* = 0.056 | *t* = 2.67<br>*p* = 0.035 | *t* =-2.44<br>*p* = 0.048 | *t* = -1.97<br>*p* = 0.12 |
| **N1** | *t* = 3.29<br>*p* = 0.007 | *t* = 0.90<br>*p* = 0.55 | *t* = 2.70<br>*p* = 0.035 | *t* = 0.99<br>*p* = 0.51 | *t* = 1.71<br>*p* = 0.2 | *t* = 1.36<br>*p* = 0.36 | *t* = -0.36<br>*p* = 0.78 | *t* = -0.08<br>*p* = 0.94 | *t* = -0.71<br>*p* = 0.61 |
| **N2** | *t* = 2.46<br>*p* = 0.048 | *t* = 0.65<br>*p* = 0.63 | *t* = 3.29<br>*p* = 0.007 | *t* = -1.22<br>*p* = 0.40 | *t* = 3.27<br>*p* = 0.007 | *t* = 1.06<br>*p* = 0.49 | *t* = -0.41<br>*p* = 0.77 | *t* = 0.76<br>*p* = 0.60 | *t* = -0.86<br>*p* = 0.55 |

**Table S5. Confusion matrix and performance scores of the random forest classifier (responsive vs non-responsive trials) in different sleep stages, for HP and NP.**

| Participants with Narcolepsy (NP) | | | | | | |
|---|---|---|---|---|---|---|
| | True positives (TP) | False positives (FP) | True negatives (TN) | False negatives (FN) | Balanced accuracy | F1 score |
| **Wake** | 671 (80.1%) | 33 (25.2%) | 98 (74.8%) | 159 (19.2%) | 0.78 (78%) | 0.87 |
| **N1** | 181 (59%) | 72 (36.4%) | 126 (63.6%) | 126 (41%) | 0.61 (61%) | 0.64 |
| **N2** | 187 (67%) | 448 (35.6%) | 810 (64.4%) | 92 (33%) | 0.66 (66%) | 0.41 |
| **REM** | 151 (65.9%) | 101 (28.2%) | 257 (71.8%) | 78 (34.1%) | 0.67 (67%) | 0.61 |
| **Healthy Participants (HP)** | | | | | | |
| | True positives (TP) | False positives (FP) | True negatives (TN) | False negatives (FN) | Balanced accuracy\| | F1 score |
| **N2** | 29 (39.7%) | 318 (25.1%) | 948 (74.9%) | 44 (60.3%) | 0.58 (58%) | 0.14 |

**Table S6. Statistical differences of the neurophysiological markers between responsive and non-responsive trials in lucid REM sleep (participants with narcolepsy).** K for Kolmogorov Complexity; PE θ for Permutation Entropy in the theta band; SE for Sample Entropy; and wSMI θ for weighted symbolic mutual information in the theta band. All p-values are FDR corrected for multiple comparisons. BF: Bayes Factor of the comparison between the full model (response + subject identity) and a null model (subject identity only).

| Lucid REM Sleep | K | PE θ | SE | wSMI θ | PSD \|γ\| | PSD \|β\| | PSD \|α\| | PSD \|θ\| | PSD \|δ\| |
|---|---|---|---|---|---|---|---|---|---|
| **Responsive vs. Non-responsive trials** | $t$ = 1.26<br>$p$ = 0.62<br>BF = 0.17 | $t$ = 0.05<br>$p$ = 0.96<br>BF = 0.08 | $t$ = 1.18<br>$p$ = 0.53<br>BF = 0.18 | $t$ = -0.58<br>$p$ = 0.75<br>BF = 0.10 | $t$ = 0.55<br>$p$ = 0.75<br>BF = 0.10 | $t$ = -1.31<br>$p$ = 0.53<br>BF = 0.21 | $t$ = -0.70<br>$p$ = 0.75<br>BF = 0.11 | $t$ = 1.20<br>$p$ = 0.53<br>BF = 0.17 | $t$ = -0.06<br>$p$ = 0.96<br>BF = 0.08 |

**Table S7. Statistical differences of the neurophysiological markers between lucid and non-lucid trials (REM sleep), for all trials and for responsive trials only.** K for Kolmogorov Complexity; PE θ for Permutation Entropy in the theta band; SE for Sample Entropy; and wSMI θ for weighted symbolic mutual information in the theta band. All p-values are FDR corrected for multiple comparisons. Statistically significant comparisons ($p$ < 0.05) are highlighted. BF: Bayes Factor of the comparison between the full model (response + subject identity) and a null model (subject identity only).

| Lucid REM sleep vs. REM sleep | K | PE θ | SE | wSMI θ | PSD \|γ\| | PSD \|β\| | PSD \|α\| | PSD \|θ\| | PSD \|δ\| |
|---|---|---|---|---|---|---|---|---|---|
| **All trials** | $t$ = 2.08<br>$p$ = 0.067 | $t$ = 2.33<br>$p$ = 0.045 | $t$ = 2.99<br>$p$ = 0.024 | $t$ = -1.13<br>$p$ = 0.33 | $t$ = 2.73<br>$p$ = 0.024 | $t$ = 1.90<br>$p$ = 0.087 | $t$ = 0.96<br>$p$ = 0.38 | $t$ = 0.72<br>$p$ = 0.47 | $t$ = -2.66<br>$p$ = 0.024 |
| **Responsive trials** | $t$ = 1.49<br>$p$ = 0.31<br>BF = 0.17 | $t$ = -0.07<br>$p$ = 0.94<br>BF = 0.14 | $t$ = 1.97<br>$p$ = 0.31<br>BF = 0.28 | $t$ = -1.62<br>$p$ = 0.31<br>BF = 0.17 | $t$ = 1.55<br>$p$ = 0.31<br>BF = 0.21 | $t$ = -0.36<br>$p$ = 0.81<br>BF = 0.19 | $t$ = -0.74<br>$p$ = 0.69<br>BF = 0.14 | $t$ = 1.37<br>$p$ = 0.31<br>BF = 0.21 | $t$ = -0.54<br>$p$ = 0.76<br>BF = 0.20 |

**Table S8. Detailed information on the sleep characteristics of the participants with narcolepsy and healthy participants.**

| Group | Nap | Wake | | N1 | | N2 | | N3 | | REM | | Sleep Latency | | Micro-arousals | | TST | | Micro-arousal Index | | WASO | | Efficiency | |
|---|---|---|---|---|---|---|---|---|---|---|---|---|---|---|---|---|---|---|---|---|---|---|---|
| | | % | std | % | std | % | std | % | std | % | std | min | std | number | std | min | std | number/H | std | min | std | % | std |
| Participants with Narcolepsy | Nap 1 | 10 | 21.21 | 16.33 | 17.85 | 24.55 | 23.7 | 2.63 | 9.44 | 46.5 | 31.57 | 2.86 | 4.65 | 11.85 | 8.37 | 15.7 | 5.51 | 52.05 | 50.56 | 1.43 | 3.32 | 90 | 21.21 |
| | Nap 2 | 4.86 | 14.65 | 25.06 | 23.46 | 40.73 | 24.7 | 7.02 | 15.17 | 21.88 | 30.34 | 3.33 | 4.63 | 9.8 | 7.76 | 16.01 | 5.16 | 36.85 | 29.16 | 0.66 | 1.67 | 95.14 | 14.65 |
| | Nap 3 | 5.05 | 7.06 | 24.74 | 23.3 | 41.03 | 24.03 | 13.33 | 22.48 | 15.85 | 28.61 | 3.48 | 4.01 | 9.19 | 7.08 | 15.77 | 4.23 | 39.73 | 35.44 | 0.75 | 0.99 | 94.95 | 7.06 |
| | Nap 4 | 8.3 | 13.93 | 18.47 | 15.75 | 42.16 | 28.44 | 11.93 | 21.01 | 19.15 | 30.76 | 2.31 | 2.44 | 10.7 | 6.74 | 16.28 | 3.59 | 42.43 | 29.5 | 1.41 | 2.49 | 91.7 | 13.93 |
| | Nap 5 | 7.58 | 13.31 | 19.49 | 15.85 | 26.04 | 22.41 | 7.91 | 21.7 | 38.97 | 34.12 | 4.22 | 4.19 | 9.79 | 7.09 | 14.73 | 4.77 | 38.67 | 32.56 | 1.05 | 1.87 | 92.42 | 13.31 |
| | Total | 7.158 | 14.032 | 20.818 | 19.242 | 34.902 | 24.656 | 8.564 | 17.96 | 28.47 | 31.08 | 3.24 | 3.984 | 10.266 | 7.408 | 15.698 | 4.652 | 41.946 | 35.444 | 1.06 | 2.068 | 92.842 | 14.032 |
| Healthy Participants | Morning N = 14 | 18.82 | 18.08 | 20.69 | 7.39 | 36.63 | 14.04 | 10.07 | 13.7 | 13.79 | 12.13 | 16.52 | 13.02 | 35.36 | 15.23 | 69.5 | 21.98 | 33.65 | 16.3 | 13.98 | 10.89 | 81.18 | 18.08 |
| | Afternoon N = 8 | 42.37 | 18.43 | 26.64 | 18.07 | 29.18 | 25.44 | 1.81 | 4.71 | 0 | 0 | 18.02 | 25.83 | 36.75 | 15.12 | 47.04 | 22.32 | 57.61 | 39.16 | 34.94 | 19.51 | 57.63 | 18.43 |
| | Total | 27.38 | 21.21 | 22.85 | 12.3 | 33.92 | 18.74 | 7.07 | 11.84 | 8.78 | 11.71 | 17.07 | 18.11 | 35.86 | 14.84 | 61.33 | 24.24 | 42.36 | 28.54 | 21.6 | 17.52 | 72.62 | 21.21 |
